# 3D-Visualization of Amyloid-β Oligomer and Fibril Interactions with Lipid Membranes by Cryo-Electron Tomography

**DOI:** 10.1101/2020.07.21.214072

**Authors:** Yao Tian, Ruina Liang, Amit Kumar, Piotr Szwedziak, John H. Viles

## Abstract

Amyloid-β (Aβ) monomers assemble into mature fibrils via a range of metastable oligomeric and protofibrillar intermediates. These Aβ assemblies have been shown to bind to lipid bilayers. This can disrupt membrane integrity and cause a loss of cellular homeostasis, that triggers a cascade of events leading to Alzheimer’s disease. However, molecular mechanisms of Aβ cytotoxicity and how the different assembly forms interact with the membrane remain enigmatic. Here we use cryo-electron tomography (cryoET) to obtain three-dimensional nano-scale images of various Aβ assembly types and their interaction with liposomes. Aβ oligomers bind extensively to the lipid vesicles, inserting and carpeting the upper-leaflet of the bilayer. Furthermore, curvilinear protofibrils also insert into the bilayer, orthogonally to the membrane surface. Aβ oligomers concentrate at the interface of vesicles and form a network of Aβ-linked liposomes. While crucially, monomeric and fibrillar Aβ have relatively little impact on the membrane. Changes to lipid membrane composition highlights a significant role for GM1-ganglioside in promoting Aβ-membrane interactions. The different effects of Aβ assembly forms observed align with the highlighted cytotoxicity reported for Aβ oligomers. The wide-scale incorporation of Aβ oligomers and curvilinear protofibrils into the lipid bilayer suggests a mechanism by which membrane integrity is lost.

## INTRODUCTION

Alzheimer’s disease (AD) accounts for more than two-thirds of dementia world-wide. A large body of evidence indicates its molecular basis centers on a small hydrophobic peptide, amyloid-β (Aβ). Cleaved from a large amyloid precursor protein the Aβ-peptide is typically 40 or 42 amino acids in length (Aβ40/42) (1). The self-association of monomeric Aβ results in a heterogeneous mixture of small oligomeric assemblies, protofibrils and amyloid fibrils which form extra-cellular plaques in the brains of AD patients. These assemblies have different biophysical and synapto-toxic properties. The interaction of Aβ with lipid membranes is believed to impede synaptic function, causing loss of cellular homeostasis which ultimately leads to hyper-phosphorylation of tau, cell death and dementia (1).

Current understanding of Aβ membrane interactions presents quite a confused picture. This may be due to different membrane systems studied in non-native conditions, and poorly defined Aβ assembly states, while different imaging and biophysical techniques employed has resulted in different aspects of the Aβ interaction being emphasized (2-4). Aβ42 oligomers have been shown to insert into cellular membranes and form large single ion-channel pores, with an internal diameter of between 1.9 and 2.5 nm (5). Alternatively, a more wide-spread carpeting of the membrane by Aβ has been proposed. This can cause a general increase in membrane conductance due to membrane thinning and the lateral spreading of lipid head-groups (6-8). Lipid extraction by Aβ oligomers from supported lipid bilayers have been imaged by atomic force microscope (AFM) and this has been likened to the effect of a detergent (9). Others have highlighted the importance of elongating fibrils at the surface of membranes, which may cause extraction and incorporation of lipid into growing Aβ42 fibrils (10). There are also studies to indicate the lipid membrane composition, in particular levels of GM1 ganglioside (11-15), and cholesterol (16, 17), can influence Aβ affinity for the bilayer. Similar effects on lipid membranes have been reported for other amyloid forming proteins, including: amylin (18-20), alpha-synuclein (21), mammalian Prion protein (22), β2Macroglobulin (β2M) (23) and Serum Amyloid A (24) which suggests a shared mechanism of membrane disruption. These behaviors draw some parallels with the toxicity mechanism described for anti-microbial peptides (4, 25).

Previously, Aβ interplay with lipid bilayers have been studied using predominantly AFM and negative-stain transmission electron microscope (TEM) (9, 11, 26-30). These techniques have the capability to reveal nanoscale details of the membrane-amyloid interaction but at the same time can be artefact-prone. AFM can only be utilized for imaging on flat and supported surfaces, such as mica. Heavy metal staining in TEM causes sample drying and structural artefacts such as flattening of spherical objects. In contrast, cryo electron tomography (cryoET) is an electron cryomicroscopy technique that can resolve unique structures in a native state, in three dimensions (3D) and at the macromolecular resolution range (31), and is particularly well suited to investigate protein/membrane systems in 3D (32), as well as amyloid fibrils (33). CryoET has recently been used to study the interaction of fibrils from the Huntingdon’s disease associated polyQ expanded protein, with membranes from cellular inclusion bodies *in situ* (34). There is also reports of cryoET studies which focus on fibrils, but not oligomers, of β2M and their interaction with liposomes (23). There has also been a room-temperature tomographic study of Serum Amyloid A fibrils stained with heavy metal (24) as well as a scanning tomographic study of Aβ plaques *ex-vivo* (35).

Here, by employing the latest developments in cryoET data collection strategies and hardware, including direct electron detectors, we report nanoscale 3D images of different Aβ assembles impacting the surface of liposomes. In contrast to monomeric and mature fibrils, oligomeric and curvilinear protofibrils interact extensively with the membrane surface, carpeting and inserting into the upper leaflet of the bilayer. The Aβ decorated membrane attracts neighboring vesicles to form a tightly zippered network of inter-connected vesicles. CryoET imaging under near native conditions reveals the mechanism by which Aβ oligomers and curvilinear protofibrils can disrupt cellular homeostasis.

## RESULTS

Using an extrusion method, we have generated large unilamellar vesicles (LUVs). This lipid membrane model has the advantage that components of the bilayer can be altered, the initial lipid composition studied includes an aqueous mixture of phosphatidylcholine (PC), cholesterol and GM1-ganglioside, with a ratio 68:30:2 by weight, buffered at pH 7.4. This lipid mixture was chosen to mimic the typical composition of the extracellular face of membranes.

A few microliters of the vesicle suspension were applied to an EM grid and vitrified in liquid ethane. This process offers the best possible structural preservation and is compatible with high-resolution imaging. Then, the sample was transferred to an electron cryomicroscope and a series of 2D images at discrete angles was acquired and computationally reconstructed into a 3D volume to produce the tomogram. The liposomes suspended in aqueous buffer, range in size, typically between 100 and 250 nm in diameter. The large unilamellar vesicles are intact and highly spherical, see Figure 1 panel a, and supplemental Figure S1. The mean lipid bilayer thickness has been measured to be 5.1±0.1 nm. Multivesicular liposomes are also observed with smaller vesicles encapsulated within the larger vesicles, as shown in Figure S1a. Consistent with this, negative-stain TEM images show vesicles largely spherical and undecorated, Figure S1b.

**Figure 1.**
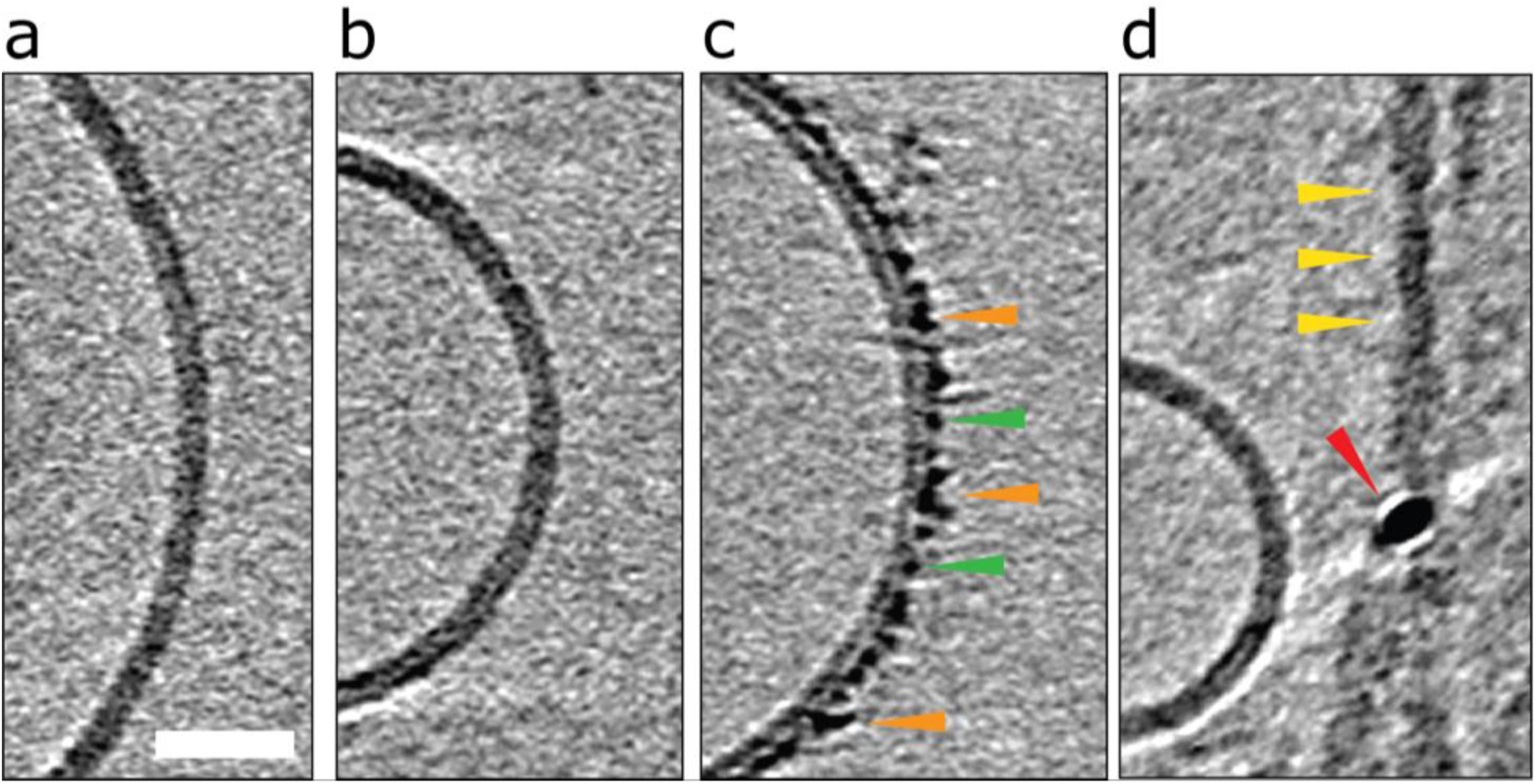
Impact of Aβ42 monomers, curvilinear protofibrils and fibrils on lipid vesicles. **a)** A typical large unilamellar vesicle in the absence of Aβ. More examples in Figure S1. **b)** Aβ42 monomer **c)** lag-phase Aβ42 oligomers/protofibrils **d)** Aβ42 fibrils. Only Aβ oligomers/protofibrils decorate the outer surface of the bilayer. Tomographic slices are 7.6 nm thick, orange, green and yellow arrowheads highlighting oligomers, curvilinear protofibrils and fibrils, respectively. The red arrowhead highlights a gold fiducial marker. Scale bar: 25 nm.

Aβ follows a nucleated polymerisation reaction in which Aβ monomers form small oligomeric assemblies that then nucleate the rapid formation of large amyloid fibrils, Figure S2. We wanted to investigate the interaction of different Aβ assembly forms with the lipid-bilayer. This was achieved by isolating and characterizing Aβ at three stages of fibril assembly. These stages were: monomeric Aβ; prefibrillar mixed oligomeric assemblies, taken from the end of the lag-phase; and also mature fibrils taken once fibril assembly has reach equilibrium, as described in the supporting information, experimental procedures.

The Aβ assemblies taken at the end of the lag-phase contain mixture of prefibrillar structures, while appreciable monomeric Aβ is still present (36), as indicated by size exclusion chromatography. Negative-stain TEM indicates the lag-phase preparations are heterogeneous and contain a number of circular oligomeric structures typically *ca.* 10 nm in diameter. When imaged by cryoET smaller oligomers with typical diameters *ca.* 2-3 nm, and many curvilinear protofibrils are observed, see Figure S3. Heterogeneous lag-phase assemblies are highly dynamic and rich in nucleating structures therefore no attempt was made to isolate these transient mixtures further.

Mature amyloid fibrils were also studied, from Aβ samples at equilibrium. These fibril assemblies were further purified by removing any smaller oligomers using a 100 kDa molecular cut-off filter. Fibrils are typically un-branched structures, 6-20 nm in diameter and microns in length, as imaged by cryoET and negative-stain TEM, shown in Figure S4.

### Aβ oligomers/protofibrils but not monomers or fibrils, decorate the liposome surface

Essentially monomeric, chromatographically purified recombinant Aβ42 was added to the liposome solution directly after elution from the size exclusion column. The final concentration of Aβ42 was 10 µM, incubated with vesicles at a concentration of 0.5 mg ml^-1^. Monomeric Aβ42 preparations were incubated with liposomes, for 10 minutes before freezing ready for cryoET imaging. Under these conditions there is some conversion of Aβ monomer to oligomers, but this is minimal. Images for lipid membranes with and without the presence of Aβ42 monomer (10 µM) shows no apparent effect, see Figure 1b and S5a. Indeed, the appearance of the lipid bilayer in the presence of monomeric Aβ42 is indistinguishable from the bilayer in aqueous buffer, Figure 1a and Figure S1, with no change in the thickness or density of the lipid bilayer.

The impact on the lipid membrane when challenged with preparations of heterogeneous oligomeric Aβ42 assemblies (10 µM, monomer equivalent) are very different compared to monomeric Aβ42. For these preparations many oligomers and protofibrils have adhered to the surface of the membrane, after 120 minutes’ incubation with the vesicles, shown in Figure 1c, additional images are shown in Figure S5b. These oligomers and curvilinear protofibril assemblies are densely adhered to the bilayer, carpeting its surface, this is also highlighted in a single threshold rendered image, Figure 2. These Aβ42 oligomers and curvilinear protofibrils have a higher density than the lipid bilayer and so from the continuation of these dense structures below the surface of the membrane, it is clear the oligomers are able to embed within the upper leaflet of the bilayer, see Figure 2c. Further examples of oligomer embedding within the membrane are shown in Figure S5d. A movie, showing Z-stacked slices through a vesicle (Supplemental Movie M1) shows the density of Aβ assemblies in the membrane, particularly at the top of the vesicle where many protruding curvilinear protofibrils and oligomers are observed above the lipid surface, this is also highlighted in supplemental Figure S6.

**Figure 2.**
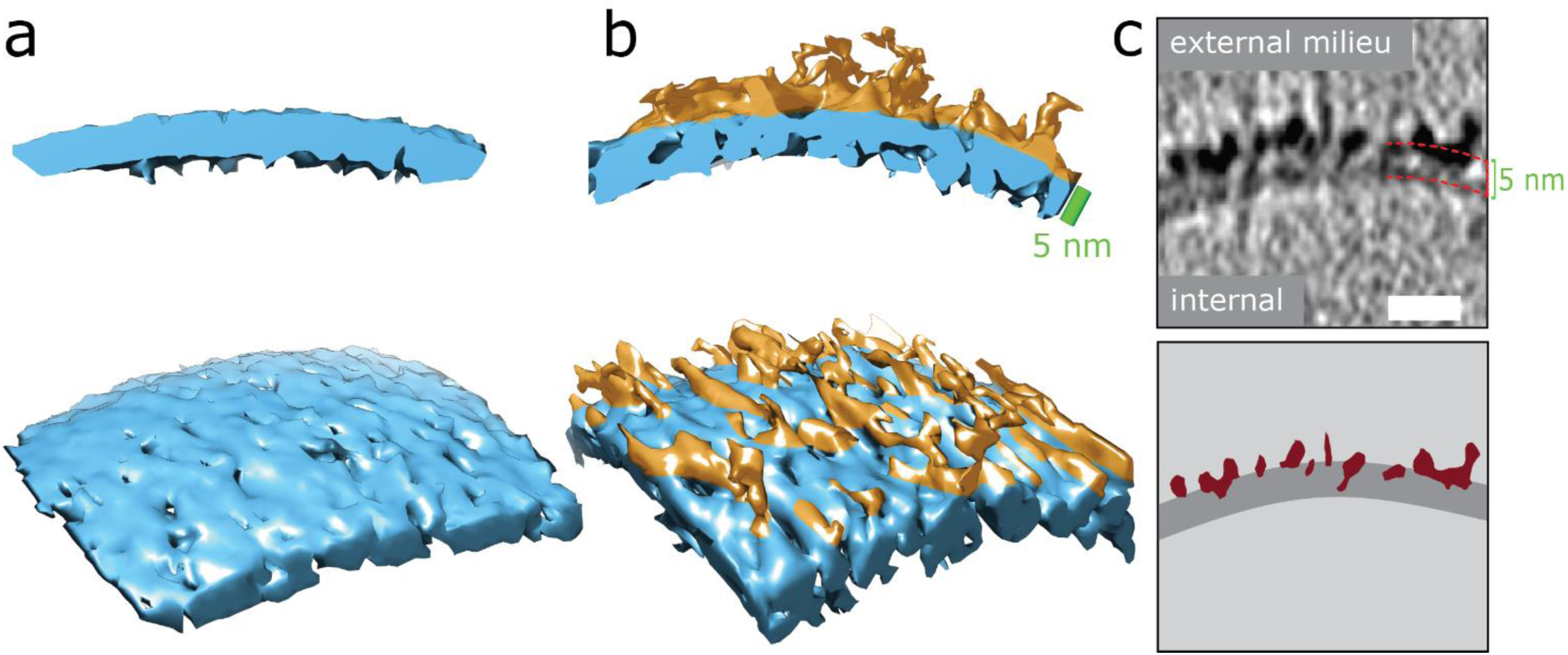
Aβ42 oligomers and protofibrils interaction with membranes. **a and b)** 3D single threshold rendered surface, image shows lipid bilayer, 5 nm thick (blue) in the absence of Aβ (a) and presence of and Aβ oligomers/protofibrils (b). Oligomers/protofibrils have carpeted the membrane. **c)** The heightened density indicates Aβ oligomer are also insert within the membrane. The same patch of membrane is also shown as a tomographic slice where the outer and inner surface are marked 5 nm apart, (top panel) and as a segmentation (bottom panel) with Aβ protofibrils and oligomers (burgundy) inserting into the lipid bilayer (dark grey), scale bar: 10 nm. Further representative images are shown in Figure S5d.

Preparations of mature Aβ42 fibrils incubated with the bilayer were also imaged, Figure 1d, and Figure S5c and S6d. The interaction of fibrils with the membrane is considerably less marked than those of the oligomeric samples. Indeed, the majority of images show no interaction between the fibrils and membrane. Unlike the oligomers, there is not an attraction of preformed fibrils to the membrane surface. The lateral face of the fibril does not readily adhere to the membrane in an aqueous environment, even when the fibril is close to the membrane, see for example, Figure S5c. Also shown in Figure S5 are the density profiles from monomer, oligomer and fibril samples, these highlight the differences in their affinity for the membrane. There are some a-typical examples, imaged by negative-stain TEM, of fibrils interacting with the membrane, these are anchored or limited to the ends of the fibrils, where fibrils do contact the membrane there are distortions and strains on the curvature of the bilayer, supplemental Figure S7. A similar behavior has been reported for β2M fibrils (23).

### Quantification indicates oligomers and protofibrils are concentrated on, and embedded within the outer-leaflet of lipid bilayers

The effects shown in Figure 1 and 2 are consistently observed for multiple vesicles as evidenced by (inspection of typically n > 50 liposomes for each condition) and multiple preparations (typically three or more for each condition). However, we wanted a way of quantifying the amount of binding and incorporation of Aβ on the lipid bilayer. Cry-oET imaging parameters included an applied defocus tuned so that the inner- and outer-leaflet of the bilayer could be resolved. Profile plots with normalized integrated intensity (NII) ‘grey-values’ were generated across the lipid bilayer. The grey values were summed for the entire 2D projection slice, 7.6 nm thick, around the whole perimeters of each vesicle, by performing radial averaging. To quantify this effect, we measured this for multiple vesicles (n=5), which equates to a summed vesicle length of typically more than 1000 nm, for each preparation. The data was collected for monomeric, oligomeric and fibrillar preparations, as well as vesicles in the absence of Aβ42. Comparisons with membrane controls in buffer alone, and with that of vesicles incubated with monomeric Aβ42 indicates no significant difference in the molecular density of the bilayer, Figure 3a, 3b and 3e. In contrast, the grey-values for the membrane incubated with Aβ42 oligomers shows considerable Aβ42 incorporation on the surface. The outer-leaflet shows a greatly increased amount of grey-value density, this indicates extensive binding on the surface and significant incorporation of Aβ42 oligomers into the membrane. Indeed, the carpeting of the bilayer has the effect of making the membrane appear thicker, as indicated in the grey-value profile plots, Figure 3c and 3e. Mean thickness of the bilayer was calculated at 5.1±0.1 nm for vesicles in only buffer, or in the presence of monomeric Aβ42 (5.1±0.2 nm) while vesicles in the presence of Aβ42 oligomers/protofibrils significantly increased the thickness by an average of 1.6 nm to 6.7±0.3 nm, Figure 3e and Figure 2 (surface rendering). This increased grey-value density is largely restricted to the outer-leaflet of the membrane, as evidenced by asymmetry between the leaflets, with the inner leaflet being on average 0.23±0.06 less dense than the outer leaflet, as shown in grey-value plot, Figure 3c and quantified in Figure 3f. Similar analysis of the grey-values across the lipid bilayer in the presence of mature Aβ42 fibrils supports the assertion that there is not a widespread interaction of Aβ42 fibrils with the membrane, Figure 3d-f, as there is no increase in the bilayer thickness and no change in the grey-value densities within the membrane.

**Figure 3.**
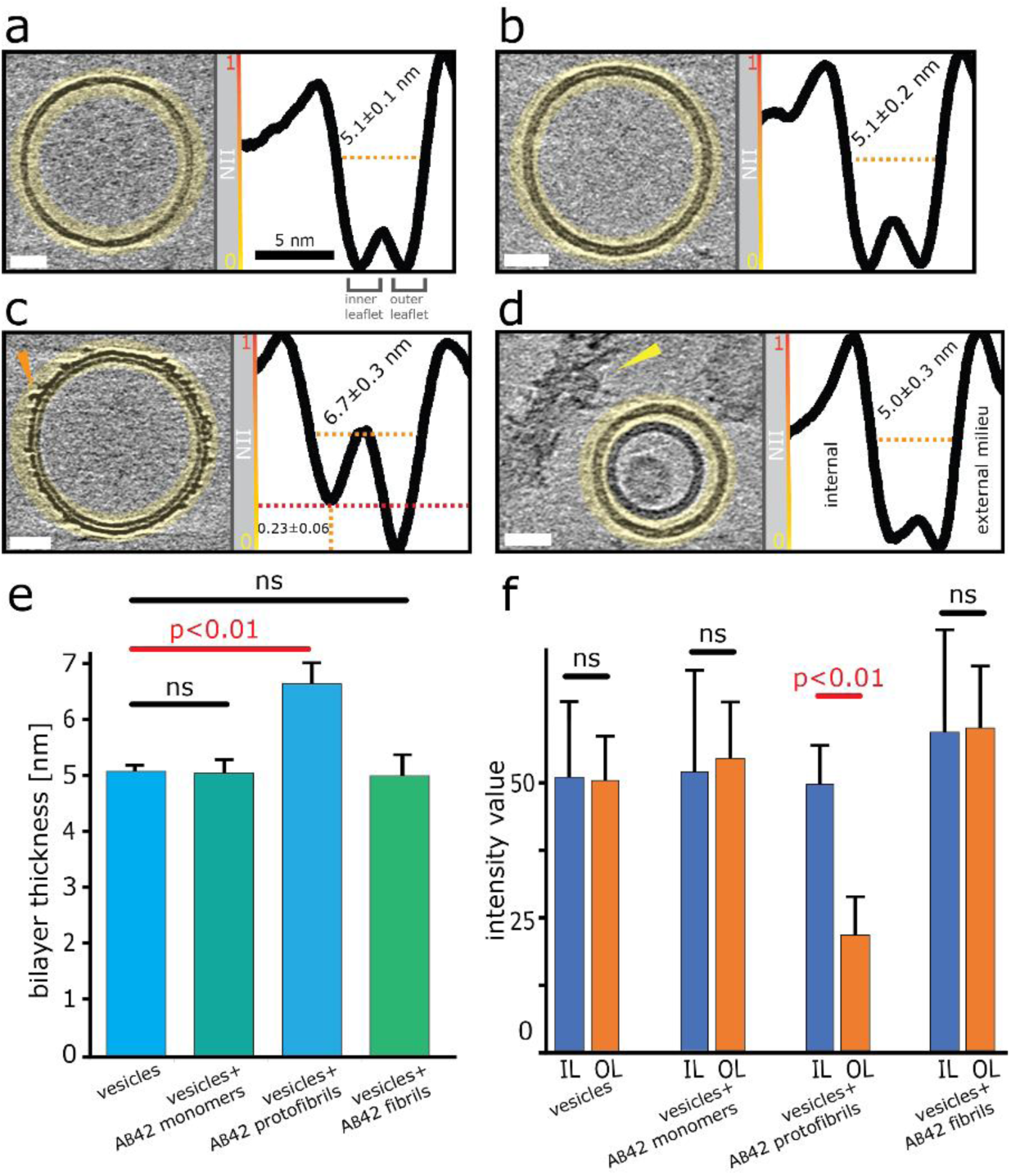
Quantification of Aβ42 insertion into outer-leaflet of lipid bilayers, measured by grey-scale intensity profile plots. a) liposome, no Aβ. b) monomeric Aβ. c) lag-phase oligomers/protofibrils. d) fibrils. Tomographic slices, 7.6 nm thick (left panels, scale bars: 25 nm) and the corresponding profile plots of normalized integrated intensities (NII) around the entire vesicle (right panels). The minimal values correspond to the dark densities; the maximum values correspond to the bright densities in tomograms areas highlighted in yellow. The double dip intensities indicate the presence of the inner- and outer-membrane leaflet. Aβ free vesicles, monomer and fibril shows similar density for the inner and outer-leaflet. In contrast for Aβ oligomers the outer leaflet has significant additional density (red dotted line). e) The thickness of the membrane, along with standard deviations, is also indicated for all four conditions. A bar-chart summarizes the thickness of the membrane measured at NII = 0.5, for 5 vesicles for each condition (>1000 nm of membrane for each condition, corresponding values are displayed on relevant panels a-d). Only Aβ oligomer/protofibrils carpet the membrane and so appear significantly thicker by an average of 1.6 nm (ANOVA analysis). **f)** The intensity values around vesicles perimeters were compared for the – inner leaflet (IL), outer leaflets (OL). Only bilayer with Aβ42 oligomers show a significant increase in density for the out-leaflet. Arrowheads point to an Aβ42 protofibril (c) and Aβ42 fibril bundle (d). See Movie M1 for Z-stacked slices through the vesicle shown in panel (c).

### 3D structure of curvilinear protofibrils embedded in lipid membranes

The numerous curvilinear protofibrils observed in the heterogeneous lag-phase preparations produce excellent high-contrast cryoET images, these three dimensional structures are represented as single-threshold surfaces, Figure 4. Corresponding structures are also shown in movies, supplemental Movie M2-M4. These assemblies have a range of lengths, see histogram in Figure S3b, the majority of curvilinear protofibrils are between 10 and 25 nm, and tend not to exceed 40 nm, with mean lengths of 19±9 nm. While their diameters are quite consistent at 2.7±0.4 nm, both values are for n=100 protofibrils. The curvilinear protofibrils varied from linear to branched structures, Figure 4a. These highly irregular and branched assemblies have considerable variation in the extent of their curvature. We believe our 3D images are some of the first curvilinear protofibrils to be imaged using cryoET. The protofibrils structures are more irregular and branched than is perhaps appreciated from 2D images and previously reported negative-stain TEM imaging.

**Figure 4.**
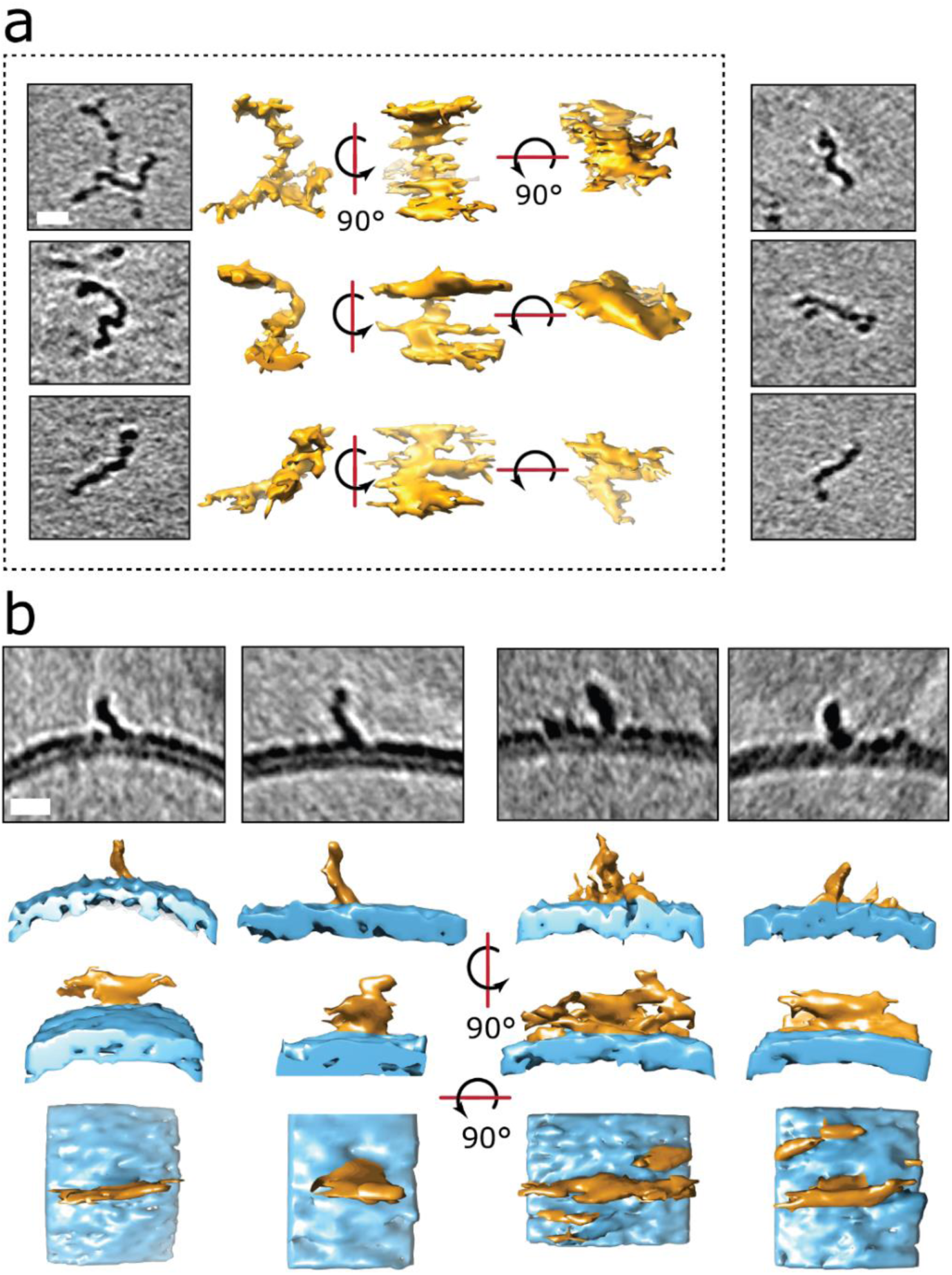
3D structures of protofibrils and their insertion into the lipid bilayer. a) Curvilinear protofibril assemblies have a variable curved appearance and can be branched. Approximately 2.7 nm diameter and up to 40 nm in length. They generate strong contrast suggesting high density of biological material, see also Figure S3. The tomographic slices are 7.6 nm thick, scale bar: 10 nm. Protofibrils presented in the dotted frame are also depicted as single threshold surfaces at three perpendicular orientations. The slightly elongated appearance of the protofibrils in the middle orientation is the effect of the missing wedge in tomography. 3D representations of the same protofibrils (dotted panel) are shown in Movie M2-4. b) Aβ42 protofibrils orthogonal to the outer leaflet of the lipid bilayer. Top panels are 7.6 nm thick tomographic slices. Lower panels are the corresponding volume data with a single threshold at three orientations. The bottom panels represent top views. Aβ42 – orange, membrane – blue. Scale bars: 10 nm.

Like the shorter oligomers these protofibrils interact with the lipid membrane extensively. There are a number of examples in which the ends of the protofibrils have embedded into the bilayer as their heightened density continues within the upper-leaflet of the bilayer, Figure 4 and S5d. The protofibrils orientated orthogonally with the membrane surface, Figure 4b, suggesting a displacement of the upper-leaflet of lipid bilayer.

It is notable that the contrast in tomographic images for the curvilinear protofibrils are more marked than that of mature fibril images under the same acquisition conditions, which is highlighted in the contrast density profile between a curvilinear protofibril, a fibril bundle and the membrane, Figure S4c. This enhanced contrast is surprising as the diameter of these protofibrils are smaller, 2.7±0.4 nm, compared to the mature fibrils, which are typically 10 nm, and range between 6-20 nm depending on the polymorph (37). This is an important observation and indicates the protofibrils have a higher density of biological material and more compact structure than mature fibrils. Interestingly, the smaller spherical oligomers have density similar to the curvilinear protofibrils, Figure S3. The diameter of these short more spherical oligomers is between 2 and 3 nm, which is similar to the diameter of curvilinear protofibrils.

### Aβ protofibrils remain on the outer leaflet and can cluster and link liposomes together at their interface

The observation that Aβ42 incorporation into the lipid bilayer is restricted to the outer leaflet, poses the question as to whether Aβ42 oligomers are able to pass through the bilayer and be observed inside vesicles. Some of the examined liposomes are multivesicular and contain a smaller vesicle encapsulated within the larger vesicle. Observation of the encapsulated vesicles, and the accompanying quantification using grey-scale analysis, indicates Aβ42 assemblies do not tend to migrate across the bilayer, as detectable Aβ42 oligomers are not observed bound to internal vesicles, Figure 5 and Figure S8 and Movie M5.

**Figure 5.**
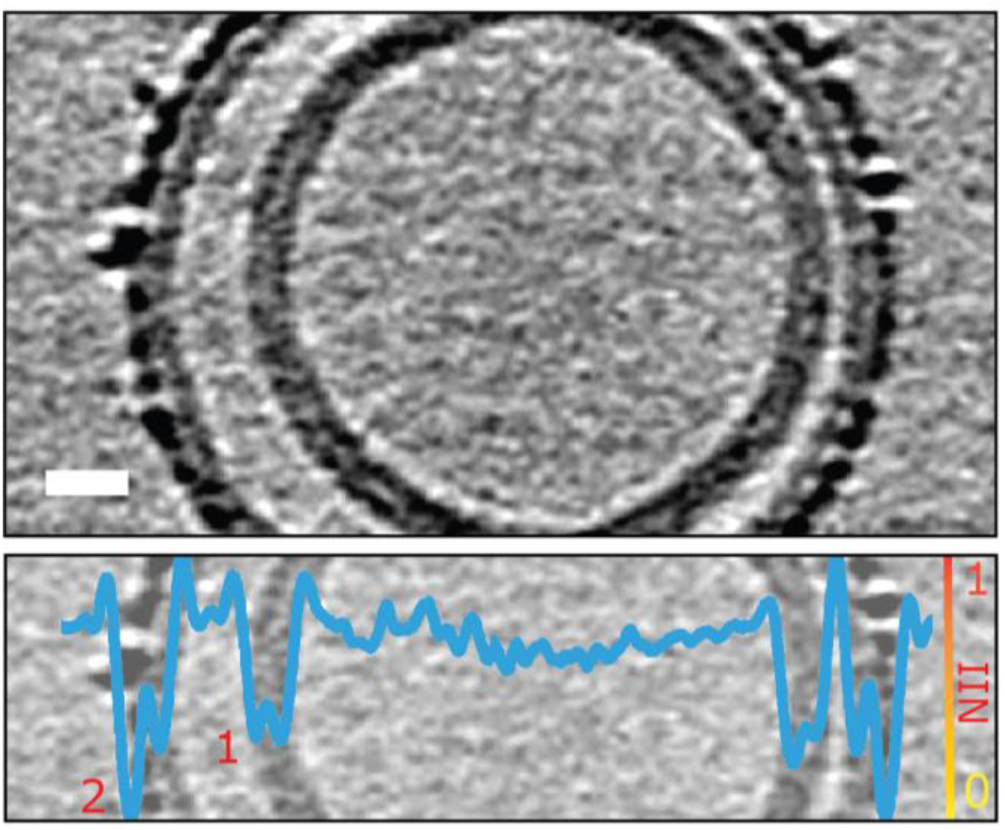
Aβ oligomers and protofibrils do not migrate to the interior of the liposome. Only the outermost liposome layer interacts with Aβ42 whereas the inner layers are protected. Shown is a multivesicular liposome (top panel) containing two lipid bilayers (numbered in bottom panel). The overlaid NII intensity plot (blue, bottom panel) indicates that the outer-leaflet of the outermost lipid bilayer (2) is densely packed with Aβ42 oligomers whereas the internal bilayer (1) is of similar density to the inner leaflet of bilayer (2) and devoid of Aβ42. The tomographic slice is 7.6 nm thick, scale bar: 10 nm. See also Figure S8 and Movie M5.

The vesicles imaged indicate that oligomers do not only have a strong attraction to the membrane surface, but they also have the effect of binding two adjacent vesicles. The presence of curvilinear protofibrils cause the contacting interface of the vesical to extend (‘zipping up’ the interface) for more than 50 nm (a typical interface of 2,000 nm^2^) causing a distortion and flattening of the vesicles at the interface. This results in the creation of a network of liposome linked together by the binding of Aβ42, as shown in Figure 6a and Figure S9. In the absence of Aβ, additional biological density is not observed between vesicles, even when the vesicles are observed contacting each other, Figure 6b. This is highlighted by the inserts showing the density between vesicles. Furthermore, in the absence of Aβ42 oligomers liposome only briefly contact each other (*ca*. 5 nm; a contact area of just 20 nm^2^) and tend not have a distortion in the spherical nature of the liposome.

**Figure 6.**
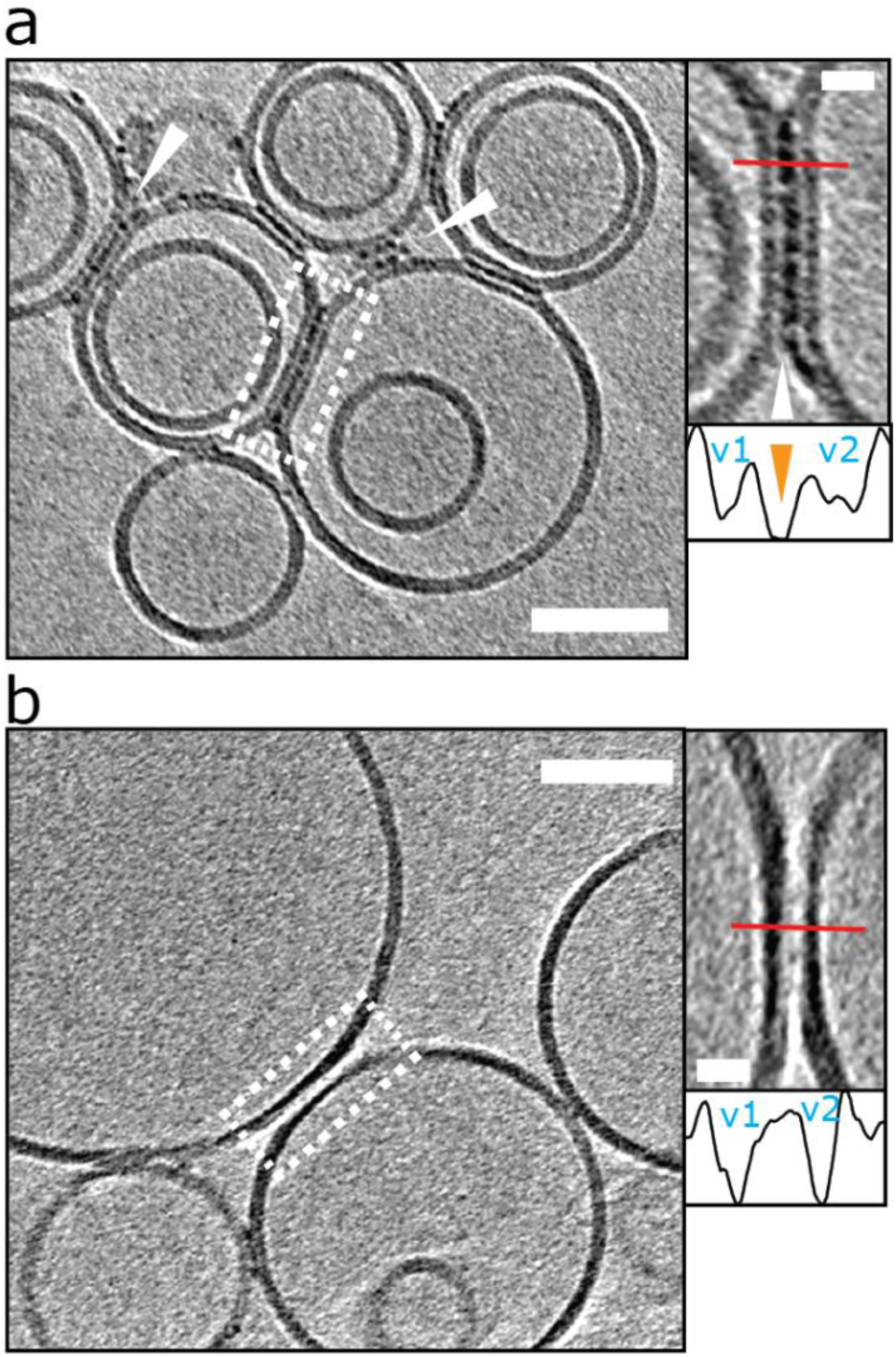
Liposomes can be linked together by Aβ42 proto-fibrils: a) For preparation with reduced levels of Aβ42 oligomer/protofibrils, Aβ42 assemblies become localized to inter-vesicular space and connect the membranes of neighboring vesicles (arrowheads). In preparations of reduced levels of oligomer/protofibrils, Aβ42 is not observed decorate elsewhere on the vesicles. Area rimmed in white is shown with more detail in inset on the right. Profile plots along the red lines indicate the presence of additional density (orange arrowhead) between the two vesicles marked as v1 and v2 b) Control experiment showing vesicles with no Aβ42 added. Please note the absence of additional densities in areas where vesicles are in contact. The tomographic slices are 7.6 nm thick with scale bars: 50 nm, insets: 10 nm. Similar supporting data is shown in Figure S9.

Interestingly this effect is more apparent when low levels of oligomers are present, typically for monomer samples that have been allowed to form a limited number of oligomers and protofibrils over a 2 hrs incubation with vesicles. The fluid nature of the bilayer facilitates the moving and clustering of the protofibrils at the interface between two vesicles. We note for the lower abundancy oligomer samples, the oligomers and protofibrils are exclusively observed at the interface between vesicles and not distributed elsewhere around the liposome, Figure 6a. In the situation of preparations with more abundant levels of oligomers (taken at the end of the lag-phase) the membrane becomes so saturated with Aβ42 oligomers, the linking between of vesicles is less widespread.

### Liposomes imaged by TEM with negatively stained samples

In addition to cryoET the same sample preparations have been imaged by negative-stain TEM, so as to compare the appearance of vesicles using this related technique. The same liposomes, but at 0.05 mg ml^-1^ have been incubated with recombinant and synthetic Aβ42 and Aβ40 monomer and oligomers (10 µM monomer equivalent). Two different heavy-metal stains were used; uranyl-acetate and phosphotungstic acid (PTA). Similar to the tomographic images, monomeric Aβ42 does not impact the appearance of the membrane, while Aβ42 oligomers causes considerable disruptions of the lipid bilayer. Particularly, for images in the presence of uranyl-acetate most vesicles exhibit a very distorted membrane surface, marked curvatures of the membrane, and the appearance of budding-off of the membrane, as shown in Figure S10. We also investigated the effect of recombinant Aβ40, there are no significant differences between the effects caused by Aβ42 and Aβ40 oligomer preparations, see Figure S10. Similar studies with synthetic Aβ preparations show the same effects on the lipid bilayer.

### Lipid bilayer composition; GM1-ganglioside is important for Aβ interaction on lipid bilayers

Next we were interested in how membrane composition might influence the extent by which Aβ42 oligomers interact with the lipid-bilayer. GM1-ganglioside has been shown to have a heightened affinity for Aβ (11-14). Phosphatidylcholine (PC) with cholesterol (70:30 by weight) vesicles were therefore produced in the absence of GM1 and incubated (120 mins) with Aβ42 oligomers. CryoET images show a large reduction in the extent to which Aβ binds to the membrane for GM1 free vesicles. This was consistently observed across many liposomes (n>50) and multiple preparations. Many vesicles remain largely free of oligomers, Figure 7a, which instead remain at the air/water interface or on the carbon grid support. Occasionally, but only for images in which vesicles are contacting each other do we observe protofibrils on the membrane. Here the protofibrils are concentrated only at the interface between vesicles, Figure 7b. The very different behavior of the same Aβ42 oligomer preparation interacting with lipid bilayer that contain GM1 (2 % by weight) is also shown, Figure 7c.

**Figure 7.**
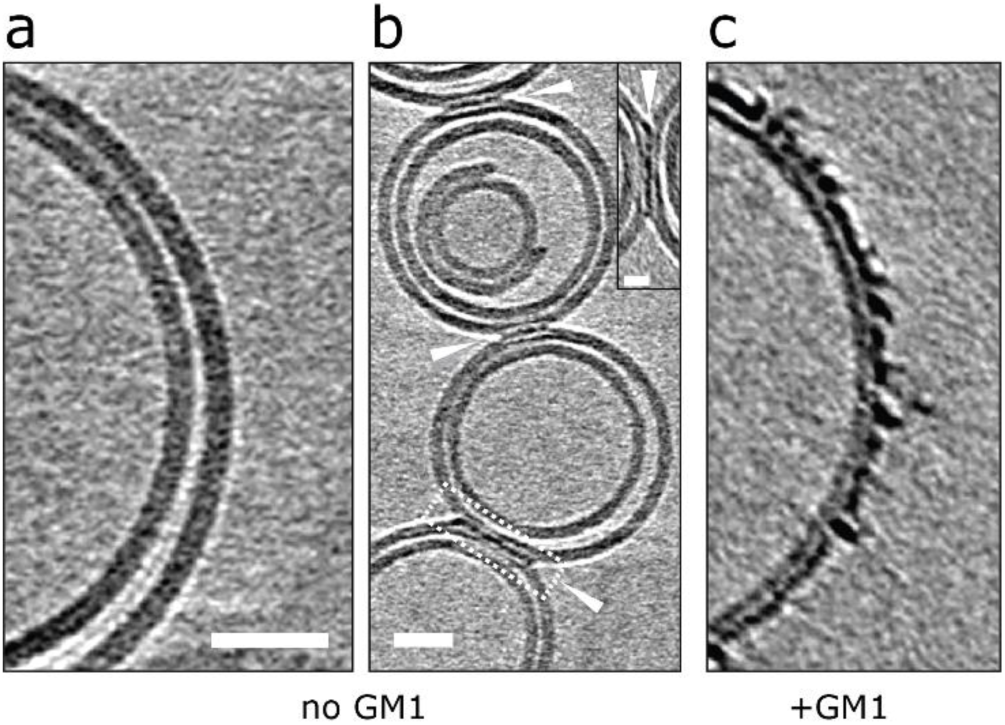
Removal of GM1-ganglioside reduces Aβ interaction on lipid bilayers. a) and b) GM1 free vesicles in the presence of Aβ42 oligomer/protofibrils (10 µM). Very few Aβ42 assemblies are observed on the lipid bilayer and are localized at the interface between vesicles but not elsewhere on the bilayer. This suggests that vesicles deprived of GM1 display a lower affinity for Aβ42. c) In contrast, GM1-containing vesicles attract Aβ42 oligomer/protofibrils efficiently. Tomographic slices are nm thick showing vesicles prepared without GM1 ganglioside and mixed with Aβ42 oligomers/protofibrils. Scale bars: 25 nm, inset: 10 nm.

A similar behavior can be observed from negative-stained samples of GM1 free vesicles, imaged by TEM. In these images there is a large reduction in the extent of membrane disruption, Figure S11. As observed for many vesicals (*ca.* 50) and multiple preparations of Aβ42 and Aβ40 oligomers. Indeed, most (90 %) of the liposomes are unaffected by Aβ42 or Aβ40 oligomers when GM1 is absent from the PC/cholesterol liposomes. This data suggests that Aβ oligomers and protofibrils display a lower affinity for GM1 deficient liposomes.

Cholesterol levels are a known risk factor in AD (38) and cholesterol has been suggested to impact interactions of Aβ with membranes (16, 17). We therefore also investigated the effect of varying the levels of cholesterol in our vesicle preparations. Phosphatidylcholine vesicles were produced with 2 % GM1 and either 9 % or 39 % (by weight) of cholesterol. These lipid mixtures also produce stable vesicles, as imaged by negative-stain TEM. Neither a reduction nor increase in cholesterol had a noticeable effect on the extent of Aβ-induced membrane disruption, supplemental Figure S12. Some images exhibit an extensive amount of budding-off of membrane from the main vesicle, imaged only when using uranyl-acetate negative-stain. In these micrographs there was also the appearance of spherical structures, the majority of which are believed to be micelles typically ca 12 nm in diameter, the larger micelles/vesicles formed are ca 20 nm in diameter, Figure S13. These, very smooth appearing, circular structures were not detected in control vesicles in the absence of Aβ, or images of Aβ oligomers alone which are less smooth and regular in appearance. Lipid extraction and small micelle formation is not apparent in our cryoET images, after incubated with monomer, oligomers or fibrils; even after 48 hrs incubation at room temperature with Aβ42 oligomers. In addition, we looked for the appearance fibril elongation nucleated at the membrane surface after 48 hrs incubation with oligomers, but this was not observed.

## DISCUSSION

We report the first 3D macromolecular description of the impact of Aβ assemblies on lipid membranes imaged by cryoET. Preparations of lipids vesicles with various Aβ assemblies in aqueous buffer produce excellent quality images in a near-native environment. In particular, the resolution achieved by cryoET makes it possible to directly distinguish impacts on the outer- and inner-leaflet of the bi-layer.

We observe very different behaviors of the various Aβ assembly forms with wide-spread insertion and carpeting of the membrane by Aβ oligomers and curvilinear protofibrils, but minimal interaction by monomers or fibrils (Figure 1-3). The carpeting of the surface of the membrane and incorporation of oligomers and protofibrils, summarized in Figure 8, will have major impact on the bilayer properties, reducing the integrity of membrane and causing leakage and the influx of ions such as Ca^2+^ in to the cell, as has been widely reported (6-8, 39, 40). The increase membrane conductance observed has been described as a thinning of the membrane, although we have shown that the insertion and carpeting of Aβ on the membrane surface actually has the effect of making the membrane appear thicker although with a reduced lipid content. The insertion of Aβ into the bilayer is localized to the outer-leaflet of the membrane. This is an important observation as it suggests that extracellular Aβ assemblies do not readily migrate to the cytosol (Figure 5). If trafficking of Aβ oligomers into the cytosol does occur *in vivo*, our data suggests it happens very slowly, or by endocytosis, or an additional membrane protein would need to assist this process.

**Figure 8.**
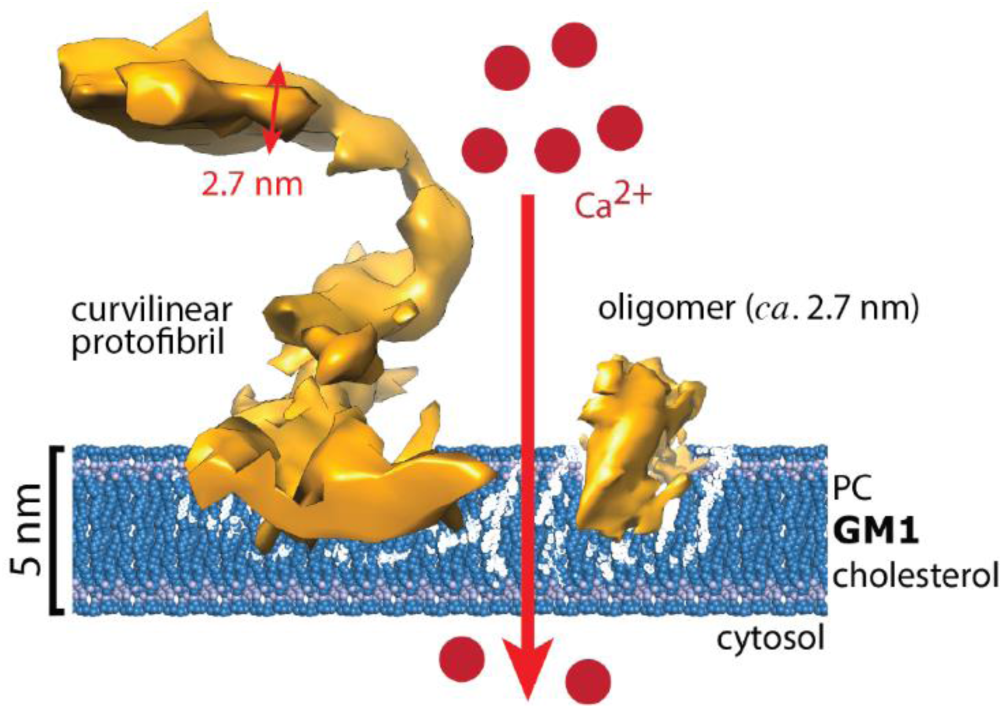
Cartoon showing Aβ42 curvilinear protofibrils and oligomers embedding in the upper leaflet of the lipid bilayer. The carpeting and incorporation of Aβ causes a general increase in permeability and conductance of the membrane and results in an influx of Ca2+ ions. The presence of GM1 ganglioside is key to the Aβ affinity for the membrane while Aβ fibrils (not shown) have minimal interaction with the bilayer.

When membranes are imaged with uranyl-acetate, as a negative-stain, the Aβ oligomers, destabilizes the membrane sufficiently to cause budding-off of vesicles and the formation of micelles (Figures S10 and S13). This effect has been likened to the action of a detergent (9, 10). Thus, we have a picture of rapid and wide-spread carpeting of membranes by Aβ oligomers and protofibrils that will destabilize the bilayer and may then lead to extraction of lipids in the long term. Indeed, Aβ amyloid plaques have a high lipid content in the AD brain (41, 42).

The high contrast density of both the curvilinear proto-fibrils facilitates the first 3D images of these assemblies. These images indicate highly irregular, branched structures, that insert into the upper-leaflet of the membrane, many of which extend out from the membrane orthogonally (Figure 4). These structures are 2.7±0.4 nm in cross-section and so could accommodate a single Aβ molecule in the one plane of the protofilament. Very similar heights, for curvilinear protofibrils have been reported using AFM, while a width of 6 nm is reported, which may reflect the lower resolution of AFM in this dimension (43). A recent AFM study of Aβ42 assemblies reports long and straight protofibrils with a regular twist, that represent the smallest of fibril structures, with a typical 5.5 nm height (44).

The higher contrast for the curvilinear assemblies and the smaller oligomers suggests a more compact structure than mature fibrils (Figure S3 and Figure S4), although fibrils are known to form tightly packed β-sheets (45, 46). Based on, in particular, the contrast density, it appears the oligomer assemblies are closely related to curvilinear protofibrils and these oligomers may simply be thought of as short curvilinear protofibrils, both of which have a diameter of *ca* 3 nm and very similar contrast density. While the grey-scale values suggest mature fibrils may be more structurally distinct (Figure S4c). Oligomers with a diameter of 3 nm suggest a protein volume ca 12-21 kDa in size. These structures can extend to form the short protofibrils which may be building blocks to longer curvilinear protofibrils (47-50). Tetrameric and octameric Aβ42 structures have been described which forms an anti-parallel β-sandwich structure in lipid bilayers (51).

The adhesive properties of Aβ curvilinear protofibrils on the surface of the lipid membrane (Figure 6) can link vesicles together. Previously unreported, the protofibrils become concentrated at the interface between two vesicles. In this way Aβ oligomers and curvilinear protofibrils might spread from one cell surface to another, by linking cells or exosomes. This suggest a mechanism by which the prion-like spreading of misfolded Aβ might occurs in the Alzheimer’s disease brain (52-54).

Our cryoET data provide a more complete picture of Aβ-membrane interactions and builds on previous studies, see reviews (2-4). The lack of interaction and impact on the membrane by monomeric Aβ is in agreement with AFM studies of supported lipid bilayers (9) and aligns with what is known about the relative cyto-toxicity of Aβ monomers and oligomers (55-57). AFM studies have reported wide-spread extraction of lipid from a mica supported lipid-bi-layer which results in the formation of large ∼50 nm holes (9, 11). This type of extraction and hole formation is not apparent in our cryoET images. The lipid within the bilayer of vesicles can behave as a fluid and fill any holes generated, if lipid extraction occurs. While in AFM studies the lipids are more immobile supported on a mica surface, which explains the different appearance observed.

Our cryoET studies in amorphous ice are well placed to study fibrils, and show minimal affinity and interaction with the membrane surface (Figures 1d, 3d, S5c and S6d).

Lateral association of fibrils on the surface of lipid membranes has been reported. However, images acquired using negative-stain and AFM are obtained by drying of samples by blotting, this causes fibrils to lay-down on the membrane surface as water is lost. Thus, for these images the interaction of fibrils may appear more widespread (9, 11, 26, 28, 29). There is a report of amyloid fibril membrane interactions, imaged by cryoET, reported for β2M fibrils (23). In this study the ends of fibrils tend to interact with the surface of the membrane. We have also observed this effect for Aβ, although the interactions with oligomers are considerably more marked. The ends of fibrils may have more exposed hydrophobic sidechains, while the lateral surface of the fibril has less of an affinity for the membrane.

There are a number of studies that describe the elevated affinity of Aβ for GM1-ganglioside compared to other lipids (11-15). We image for the first time the very different impact Aβ oligomers have on lipid membranes which do not contain GM1 (Figures 7 and S11). Aβ is negatively charged, at neutral pH, and electro-static attraction to the polar carbohydrate groups may promote the Aβ-bilayer interaction. This effect may be important as GM1-ganglioside is particularly abundant in the outer-leaflet of neuronal plasma-membranes (58).

In conclusion, 3D nanoscale imaging of liposomes suspended in a near-native aqueous environment have revealed new details and insight in to the interaction of Aβ assemblies with lipid membranes, Figure 8. Wide-scale impacts on the membrane are restricted to oligomeric and curvilinear protofibrillar structures that saturate the outer-leaflet of the membrane, while in the case of isolated monomers, or even fibrillar Aβ42, the lipid bilayer remains relatively unperturbed. This carpeting and insertion has previously been shown to impact membrane integrity and cellular homeostasis (6-8, 39, 40), and is in line with the relative cytotoxicity of Aβ oligomers compared to fibrillar assembly states (55-57). The conclusions drawn here may have many parallels for anti-microbial peptides (59, 60), and other amyloid forming proteins such as: amylin (18-20), alpha-synuclein (21), mammalian Prion protein (22), β2M (23) and Serum Amyloid A (24). Therapeutic molecules, that block insertion of Aβ oligomers into membranes may help maintain neuronal homeostasis and slow the cascade of events that culminates in dementia.

## Supporting information

MovieM1

MovieM2

MovieM3

MovieM4

MovieM5

## ASSOCIATED CONTENT

**Supporting Information** includes the experimental details and supplemental Figures S1-S13. In addition, five tomographic movies are available M1-M5.

## AUTHOR INFORMATION

### Author Contributions

All authors have given approval to the final version of the manuscript.

## ABBREVIATIONS

3D: three dimensions;
Aβ: amyloid β;
AD: Alzheimer’s disease;
AFM: atomic force microscopy;
GM1: monosialotetrahexosl-ganglioside;
CryoET: cryo-electron tomography;
Gua: HCl guanidine-hydrochloride;
HEPES: 4-(2-hydroxyethyl)-1-piperazine-ethanesulfonic acid;
IPTG: isopropyl thio-β-D-galactosidase;
NII: normalized integrated intensity;
PC: egg phosphatidylcholine;
PTA: phosphotungstic acid;
SEC: size exclusion chromatography;
TEM: transmission electron microscopy;
ThT: thioflavin T;
Tris: tris(hydroxymethyl)aminomethane;
β2M: β2Macroglobulin

