## Supplementary material for "3D-Visualization of Amyloid-β Oligomer and Fibril Interactions with Lipid Membranes by Cryo-Electron Tomography": Methods and Supplemental Figures 1-13

**Materials and Methods:**

**Supplemental Figures: Figures S1- S13**

**Movies:**

**Movie M1**: Tomogram of the vesicle shown in Figure 3C. Scale bar: 25 nm.

**Movie M2-M4**: Single threshold surfaces showing 3D representations of Aβ42 protofibrils shown in the rimmed area of Figure 4A. Scale bar: 10 nm.

**Movie M5**: Tomogram of the vesicle shown in the top left panel of supplemental Figure S8. Scale bars: 50 nm.

### Materials and Methods

#### Aβ recombinant expression

The expression and purification of recombinant Aβ40 and Aβ42 peptide was carried out using the protocol described by Walsh et al (61). Briefly, the MAβ(40/42)pETSac plasmid of Aβ40 or Aβ42 were transformed into E. coli BL21 (DE3) cells. The protein expression was induced by 1.0 mM isopropyl thio-β-D-galactosidase (IPTG) at an OD600nm of 0.5-0.6. After 4 hrs induction, cell cultures were centrifuge for 15 min at 9000 g at 4 °C, and resuspended in buffer with 10 mM Tris-Cl, 1 mM EDTA at pH 8.5. Cells were then sonicated for 30 s for one cycle at 22% amplitude (4 W). The suspension was centrifuged in 15 min at 25,000 g at 4 °C. The sonication and centrifugation steps were repeated once. The resulting pellet, containing Aβ, was dissolved in 12.5 mL denaturant buffer (8 M urea, 10 mM Tris.Cl, 1 mM EDTA at pH 8.5) with stirring overnight at 4 °C. The suspension was sonicated for 30 s and centrifuge for 15 min at 25,000 g at 4 °C to remove insoluble material. The supernatant which contains solubilized Aβ40 or Aβ42 peptide was retained.

The Aβ protein was purified using DEAE-Sepharose ion exchange resin. The peptide was eluted five steps with the same elution buffer (10 mM Tris.Cl, 1 mM EDTA, 125 mM NaCl and pH 8.5). Eluted protein was dialyzed in 20 mM ammonium bicarbonate buffer and subsequently lyophilization was carried out. The lyophilized Aβ was resolublized in 50 mM Tris.Cl, 7 M guanidine-hydrochloride (Gua-HCl) at pH 8.5 (5 ml). Solubilized peptide was further purified with size exclusion chromatography (SEC) on a Superdex 75 16/600 column (GE Healthcare) using either assay buffer (30 mM HEPES, 160 mM NaCl, pH 7.4) or 20 mM ammonium bicarbonate buffer. Aβ was used directly from the SEC elusion or lyophilized (ammonium bicarbonate buffer). The purity of Aβ was verified using 4-20% gradient SDS-PAGE. Aggregation property was assessed by thioflavin-T assay and negative stain electron microscopy.

#### Synthetic Aβ Peptides

Synthetic Aβ40 and Aβ42 was purchased from EZBiolab Inc in a lyophilized form. The majority of the data shown is for recombinant Aβ42 while images for Aβ42 fibrils were for both recombinant (Figure S6) but also for synthetic Aβ42 fibrils (Figures 2d and S5c). There was no difference in the appearance of vesicles incubated with Aβ assemblies from a recombinant or a synthetic source.

#### Monomeric Aβ by Size-Exclusion Chromatography (SEC)

The purified lyophilized Aβ40 and Aβ42 peptides were solubilized at 0.7 mg $ml^{-1}$ in water at pH 10. The solution was then placed on a shaker plate with gently rocked for 2 hours at 4°C and stored at -80°C. Monomeric Aβ was isolated using SEC with a Superdex 75 10/300 GL column (GE Healthcare). The column was pre-equilibrated with 4-(2-hydroxyethyl)-1-piperazine-ethanesulfonic acid (HEPES) (30 mM) and 160 mM NaCl buffer at pH 7.4. The flow rate of 0.5 ml $min^{-1}$ was used. SDS-PAGE was used for analyzing the collected fractions for the presence of the monomeric Aβ. The Aβ40 and Aβ42 concentration were determined using UV absorbance at 280 nm ($\varepsilon$ = 1280 $M^{-1}$ $cm^{-1}$). Monomeric samples were immediately stored after SEC elution at -80 °C. Negative-stain TEM and ThT fluorescence confirmed that SEC-purified Aβ was seed-free. Thioflavin T (ThT) fluorescence showed SEC-purified Aβ had no ThT fluorescence signal and exhibited a clear lag-phase, shown in Supplemental Figure S2.

#### Aβ Oligomer and Fibril Preparations

Aβ40 and Aβ42 monomer (10 μM) were placed in in a 96-well plate in NaCl (160 mM) and HEPES (30 mM) buffer at pH 7.4 as previously described (62). Fibril growth kinetics were monitored using a fibril-specific fluorescent dye, thioflavin T (ThT) (20 μM). Experiments were recorded using a Fluostar Omega fluorescent plate reader (BMG Labtech, Aylesbury, UK), with an excitation filter at 440 nm and an emission filter at 490 nm. Fluorescence reading at every 30 minute intervals, following well plates were agitated for 30 seconds. Adjacent sample-wells with no ThT added were used in all experiments.

Aβ40 and Aβ42 prefibrillar assemblies with predominantly oligomeric and curvilinear protofibril structures, were obtained from the well plate towards the end of the lag-phase, as monitored by ThT fluorescent dye in separate wells. Oligomeric samples were used immediately or stored at -80 oC to halt further assembly. Lag-phase mixed prefibrillar assembles were characterize by cryoET and TEM, supplement Figure S3.

At equilibrium (as indicated by ThT fluorescence in separate wells) Aβ assemblies had the typical amyloid fibrous appearance according to TEM, see Supplemental Figure S4. Fibril preparations were centrifuged using 100 kDa molecular cut-off filter (Amicon Ultra) to remove any low molecular weight oligomers. Fibril preparations have a low level of Aβ monomer and oligomer content.

**Vesicle Preparation**

Large unilamellar vesicles (LUVs) were produced using an extrusion method described previously (29). The lipids used were egg phosphatidylcholine (PC) dissolved in chloroform (Avanti Polar Lipids inc.); monosialotetrahexoslganglioside (GM1) dissolved in ethanol: H2O (1:1) (Avanti Polar Lipids inc.); and cholesterol dissolved in chloroform (Sigma-Aldrich Company Ltd.). Most of the studies used a mixture of 68:30:2 by weight of PC: chloroform: GM1. Lipid solutions were placed in a fume hood overnight to allow chloroform evaporation. Lipids film were then re-solubilized in NaCl (160 mM) HEPES (30 mM) buffered at pH 7.4, to a lipid concentration of 1 mg $ml^{-1}$. LUVs were formed from solubilized lipids using a benchtop mini extruder (Avanti Polar Lipids, Alabama, USA). The lipid solution was passed across a polycarbonate membrane with 100 nm pores 21 times. Vesicles were then stored at 4°C until use, for up to 2 days and characterized by cryoET, Supplemental Figure S1. Vesicles with elevated or reduced cholesterol were also produced, in particular PC:cholesterol:GM1 in varying ratios (89:9:2 and 59:39:2 by weight). In addition, vesicles were generated in the absence of GM1 (PC: cholesterol, 70:30 by weight). Initial studies confirmed freezing vesicles in aqueous buffer had no apparent effect on the morphology according to negative-stain TEM. Unless otherwise stated all other chemicals were purchased from Sigma-Aldrich.

**Cryo electron tomography (cryoET) and Image processing**

The final lipid vesical concentration was 0.5 mg $ml^{-1}$ for cryoET imaging with recombinant Aβ42 (5 μM monomer equivalent). Aβ42 was incubated with the vesicles for 10 and 120 mins for Aβ42 monomer preparations; 120 min and 48 hrs for lag-phase oligomers preparations; and 120 mins for Aβ42 fibrils before plunge-freezing. Vesicle solutions were plunge-frozen onto Quantifoil R2/2 holey carbon grids using a Thermo Fisher Vitrobot.

Electron cryo-tomography was performed using a Thermo Fisher Glacios TEM operating at 200 kV, equipped with a 4k x 4k Falcon 3EC direct electron detection camera at a magnification of 73k, corresponding to a pixel size of 1.9 Å at the specimen level. Specimens were tilted from approximately –60° to +60 with a 3° increment using the dose symmetric scheme. The defocus was set between 3 and 4 μm, and the total dose for each tilt series was approximately 100 e/$Å^2$. Final tomograms were binned 4x, with a pixel size of 7.6 Å. Tomographic slices were typically shown as an average of 10 slices, 7.6 nm thick.

Tomographic reconstructions from tilt series were calculated using RAPTOR (63) and the IMOD tomography reconstruction package, followed by SIRT reconstruction with the TOMO3D package (64, 65). Measurements of distances between structures were carried out within IMOD, the length of protofibrils followed the curvature of the protofibrils in three dimensions manually plotting through the center of mass using the 3dmod model modality. Single threshold surface representations and movies were prepared using Chimera (66), using a similar threshold level to exclude most background noise for each data set. The choice of the threshold level is subjective and depends on the density distribution and background artifacts.

Analysis presented in Figure 2, grey-values were measure all around perimeters of vesicles by performing radial averaging using the extended Radial Profile plugin of ImageJ (67). Profile plots of normalized integrated intensities around concentric circles as a function of distance from a point in the center of each 2D projection of a vesicle were generated using ImageJ.

**Negative-stain TEM**

Vesicles (0.05 mg $ml^{-1}$) were imaged by neg-stain TEM in the absence and presence of Aβ (10 μM monomer-equivalent) incubated for 120 mins. 5 μl aliquots of sample were added to glow discharged carbon-coated 300-mesh grids (Agar Scientific Ltd) by the droplet method, then blotted after 90 seconds and rinsed with ddH20. Following this, (5 μl) uranyl acetate (2% g/100 ml) was added, then blotted and rinsed after 10 seconds. Vesicle samples were also stained with phosphotungstic acid (PTA) (2% g/100ml) data not shown. Images were at 80,000 magnification by a JEOL model JEM-1230 electron microscope (JEOL, Ltd., Japan), operated at 80 kV, paired with a 2k Morada CCD camera and corresponding iTEM software (Olympus Europa, UK).

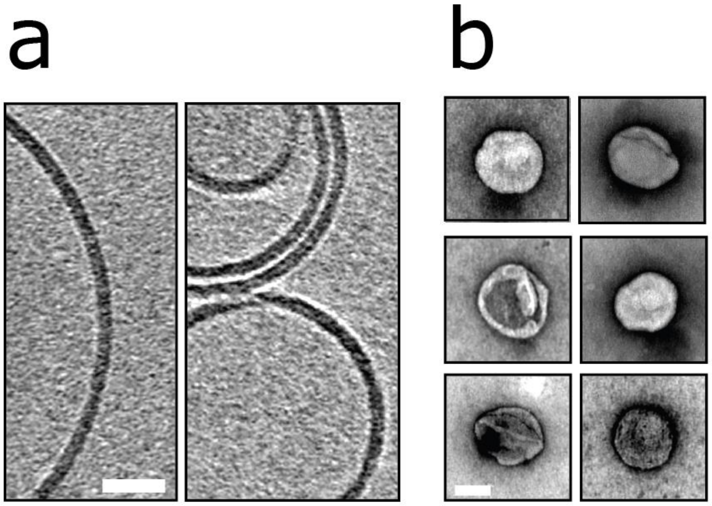


**Figure S1: Lipid Vesicles.** Liposomes were generated by the extrusion method using a lipid mixture of PC: Cholesterol: GM1, with a lipid of ratio 68: 30: 2 by weight. **a)** CryoET indicates large unilamellar vesicles, typically 100-250 nm in diameter are produced by this method. These are spherical and have a smooth regular appearance where the inner and outer leaflets of the lipid bilayer are indistinguishable by their appearance. There are also examples of multivesicular and occasionally multilamellar liposomes. Tomographic slices are 7.6 nm thick, scale bar: 25 nm. **b)** Vesicles are imaged by negative-stain TEM for the same vesicle preparation, stained with uranyl-acetate. The vesical remain largely circular and intact. These vesicles have a deflated appearance due to the drying effect of negative-stain, scale bar 100 nm.

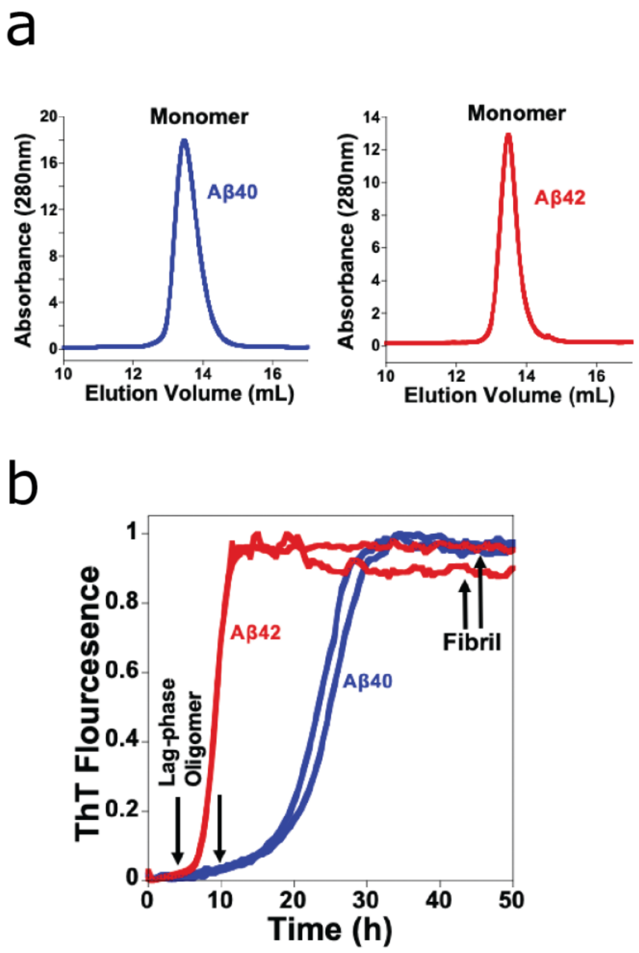


**Figure S2: Isolation of Aβ monomer, oligomer and fibril. a)** Solubilised recombinant and synthetic Aβ40, and Aβ42 was purified by elution through size-exclusion column (Superdex 75 10/300 GL column). The elution profile (280 nm) indicates a single monomeric fraction of recombinant Aβ40 and Aβ42. The Aβ monomeric samples were taken directly from the SEC column. **b)** ThT fluorescence fibre growth assays for recombinant Aβ40 and Aβ42 (10 µM) in aqueous buffer containing NaCl (160 mM), HEPES (30 mM), at pH 7.4. Aβ samples (in which ThT had not been added) were taken from the same well-plate at the appropriate time-points. Samples designated oligomeric Aβ were taken from wells towards the end of the lag-phase while fibril samples were taken from the well plate once ThT fluorescence signal had plateaued.

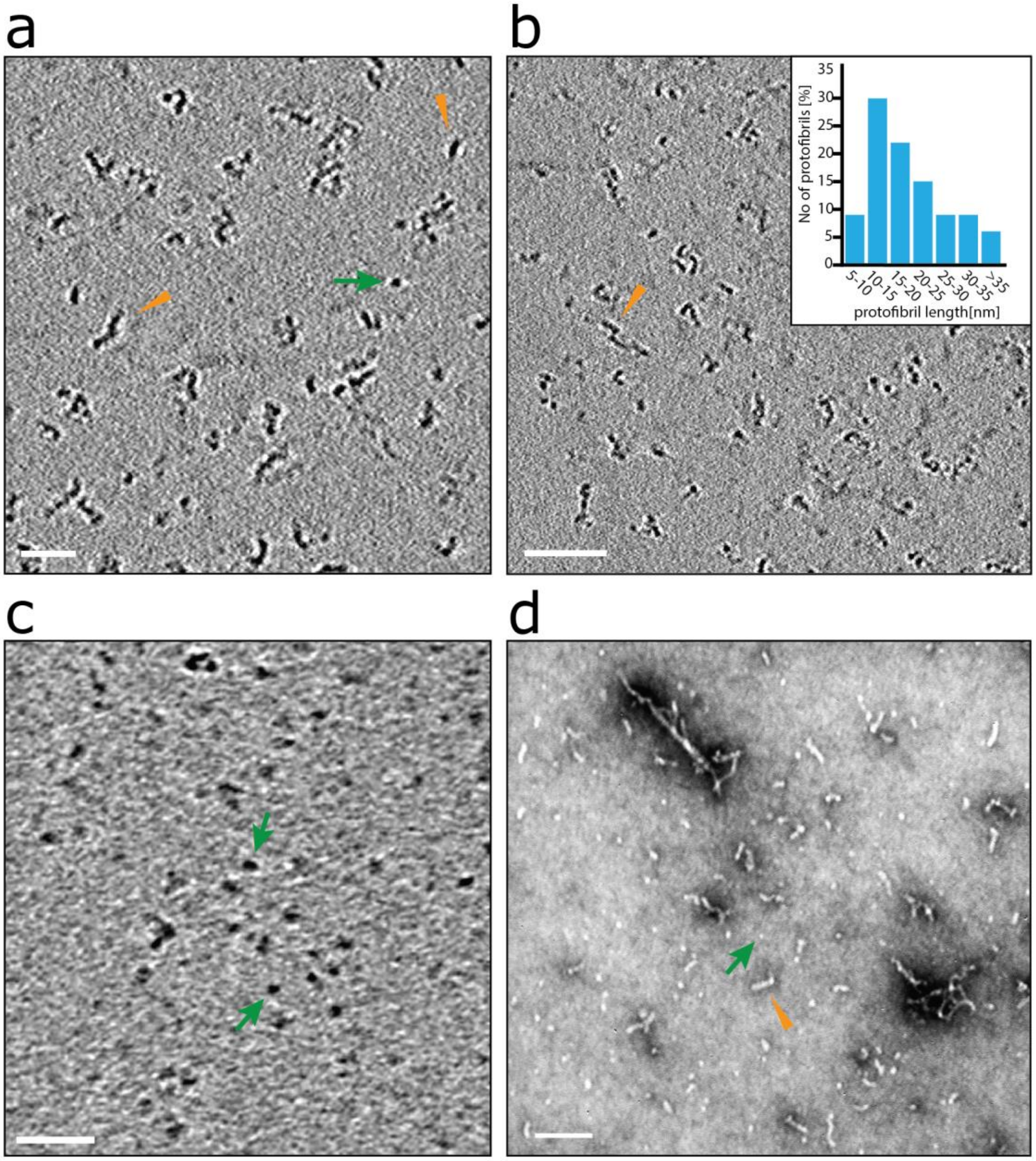


**Figure S3: Lag-phase Aβ42 oligomers and curvilinear protofibrils, imaged by cryoET and negative-stain TEM. a) and b)** The range of curvilinear protofibrils observed in a tomographic slice (7.6 nm thick) showing Aβ42 protofibrils assemblies (orange arrowheads). The assemblies are very variable in their curvature and can be branched. These structures are typically observed on the carbon support and air/water interface. Histogram shows the range of protofibril lengths (n=100), the majority of curvilinear protofibrils are between 10 and 25 nm, and tend not to exceed 40 nm. Protofibrils diameters are consistently measured to be 2.7±0.4 nm. **c)** Also observed are shorter protofibrils, described as oligomers (green arrows), approximately 3 nm in diameter but can be longer as they become curvilinear protofibrils. The spherical oligomers of *ca* 3 nm diameter suggest a molecular weight of between 12-21 kDa. Using the relationship: Volume ($nm^3$) = Mass (kDa) * 1.27 ($nm^3$/kDa). **d)** The same preparations are imaged by negative-stain (uranyl-acetate) showing curvilinear protofibrils and oligomeric structures. Note at 4.2 kDa Aβ monomer is too small to be visualized by EM. Scale bars are: a=25 nm; b=50 nm; c=25 nm; d=50 nm.

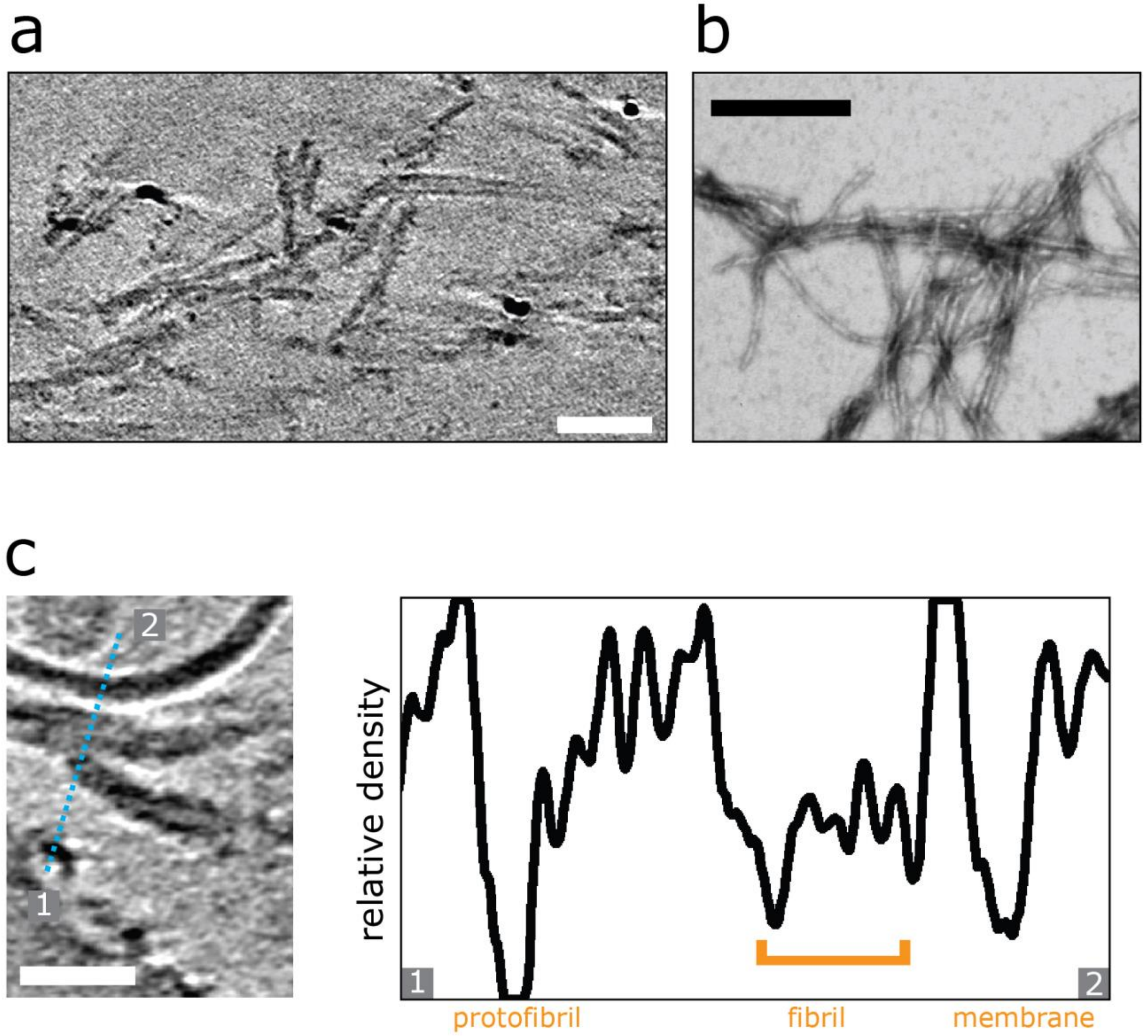


**Figure S4: Aβ42 fibrils imaged by cryoET and negative-stain TEM. a)** Tomographic slice (7.6 nm thick) for Aβ42 fibrils (10 µM). It is notable the image contrast for fibrils is less than that for the oligomers and curvilinear protofibrils. Scale bar: 50 nm. **b)** TEM negative-stain (uranyl-acetate) image of Aβ42 fibrils. Fibrils are typical 10 nm in diameter and many 100's nm in length, structures are unbranched. Scale bar: 200 nm. **c)** Tomographic slice showing Aβ42 protofibrils and fibrils (left panel) along with the respective density plot (right panel) indicating that the Aβ42 protofibril species have greater density that the fibrils. Scale bar: 25 nm.

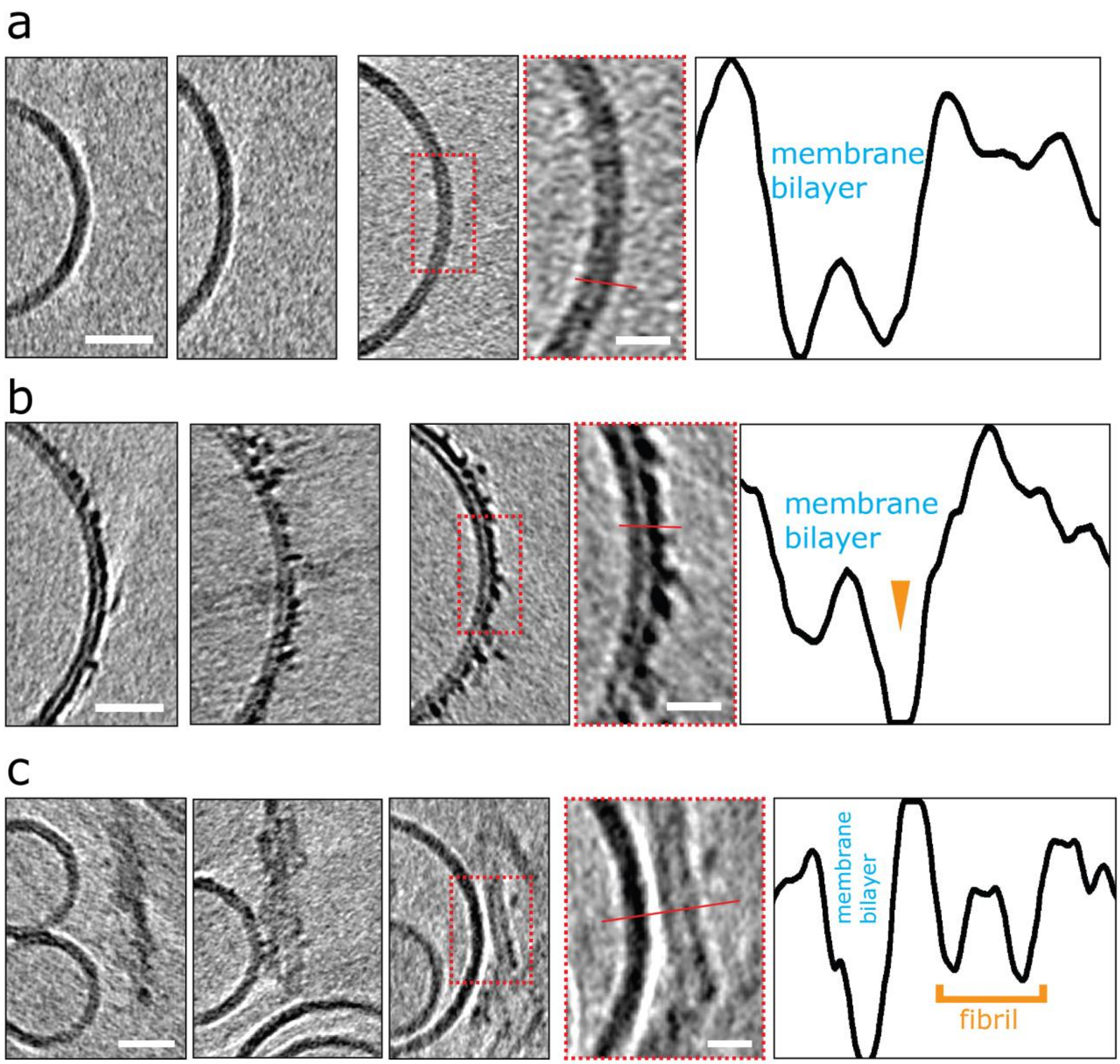
a
membrane
bilayer
b
membrane
bilayer
c
membrane
bilayer
fibril

d

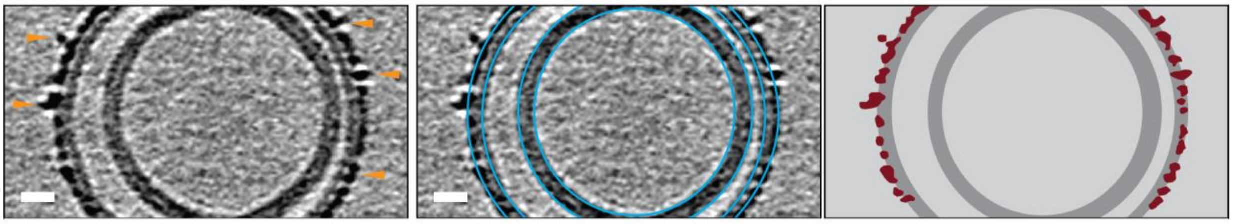

**Figure S5**: **CryoET images compare the impact of Aβ42 monomers, protofibrils and fibrils on lipid vesicles.** Tomographic slices (7.6 nm thick) showing **a)** monomer **b)** lag-phase oligomers/protofibrils **c)** fibrils. Only Aβ oligomers and curvilinear protofibrils decorate the outer surface of the bilayer. Scale bars: 25 nm. Areas rimmed in red are presented with more detail in the right outermost panels. Scale bars: 10 nm. The insets highlight the increased density on the outer leaflet of the oligomer preparation (b), while for the fibril preparation there is a lack of density even though the lateral face of the fibril aligns closely with the surface of the membrane (c). Profile plots (right panels) along the red lines indicate the asymmetry of the lipid bilayer density in (b) indicating that the outer leaflet is populated with Aβ oligomers and protofibrils (orange arrowhead). The density corresponding to a fibril in (c) is at a distance from the lipid bilayer indicating no membrane association. **d)** A tomographic slice where Aβ oligomers and protofibrils embedded in the lipid bilayer are marked with orange arrowheads (left panel), middle panel represents membranes highlighted with blue circles, and the right panel shows a segmentation with Aβ protofibrils (burgundy) inserting into the lipid bilayer (dark grey). Please note that the inner vesicle is not decorated with Aβ. Scale bar: 10 nm.

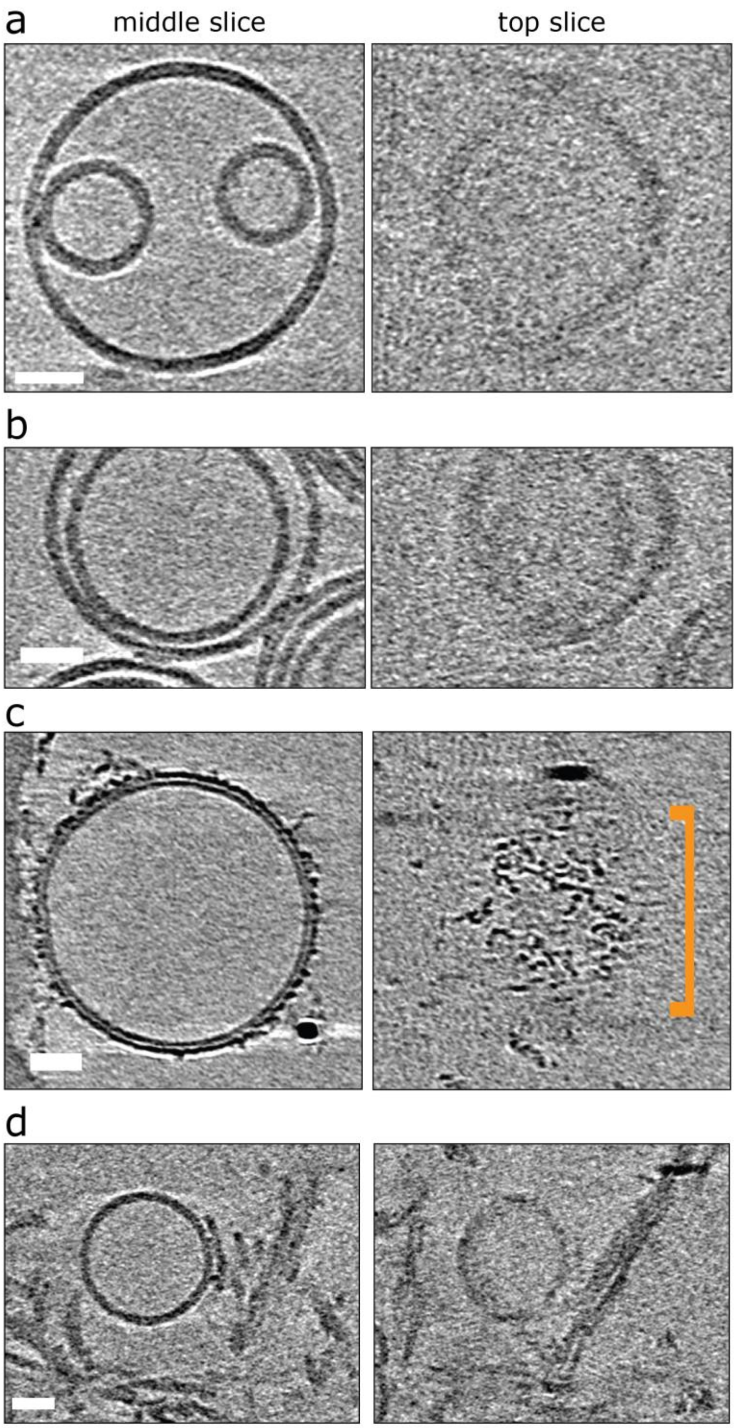


**Figure S6**: **Top and middle slices of cryoET images compare the impact of Aβ42 monomers, protofibrils and fibrils on lipid vesicles.** Tomographic slices (7.6 nm thick) showing vesicles mixed with various Aβ42 species. Left panels are middle sections, right panels are top sections. **a)** no Aβ **b)** Aβ42 monomer **c)** lag-phase oligomers/protofibrils **d)** fibrils. Please note that only vesicles in panel (c) are decorated with Aβ42 (orange marker). Scale bars: 25 nm.

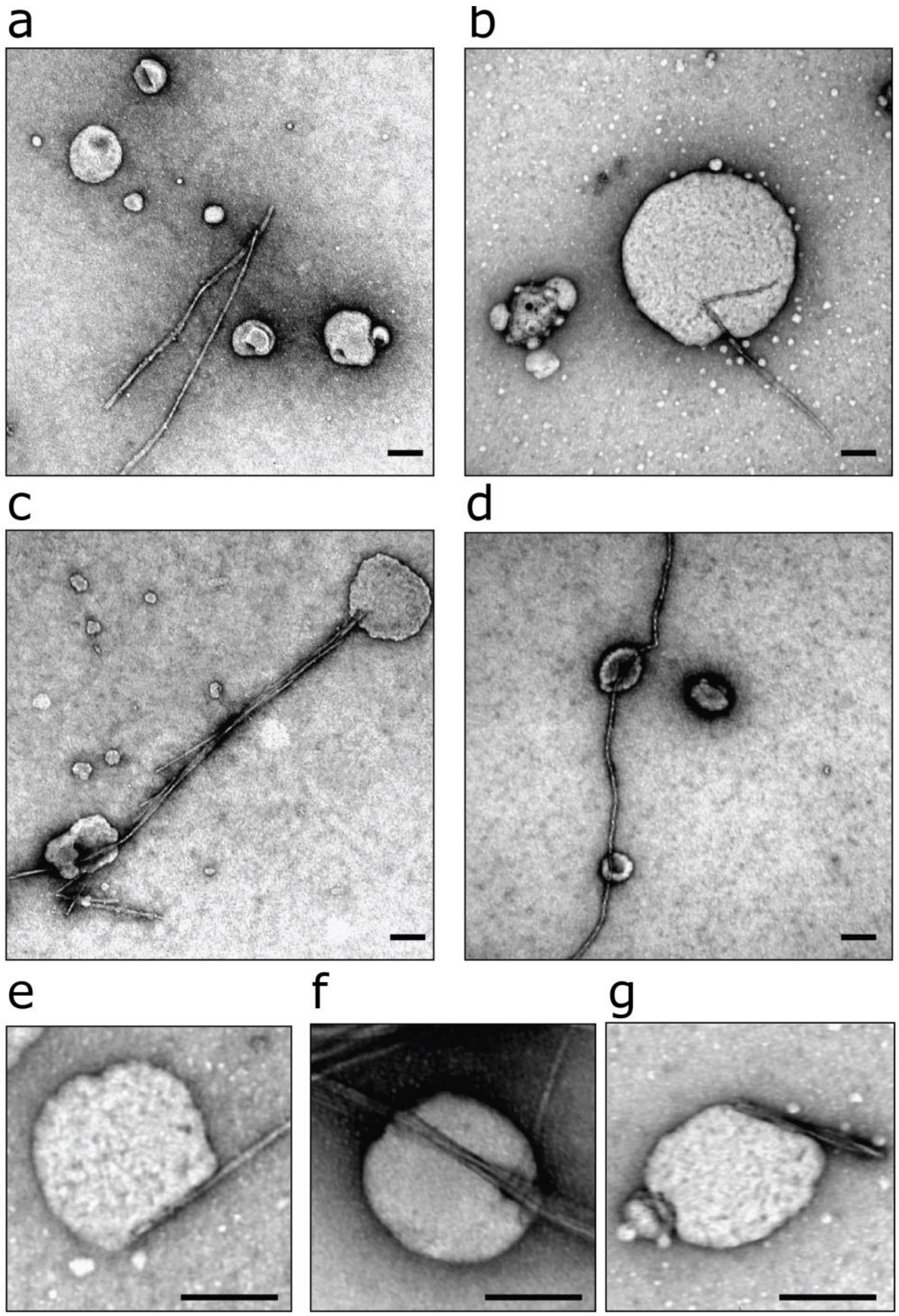


**Figure S7**: **Negative-stain TEM of vesicles with Ab42 fibrils. a)** Typical negative-stain (uranyl-acetate) image shown; the lateral face of the fibril does not readily adhere to the membrane. There are also some less common examples of fibrils interacting with membrane, these are anchored or restricted to the ends of the fibrils **(b-g)**. Lipid vesicles contain, PC:Cholesterol:GM1 (68:30:2, by weight) 0.05 mg $ml^{-1}$, pH 7.4. Scale bar: 100 nm.

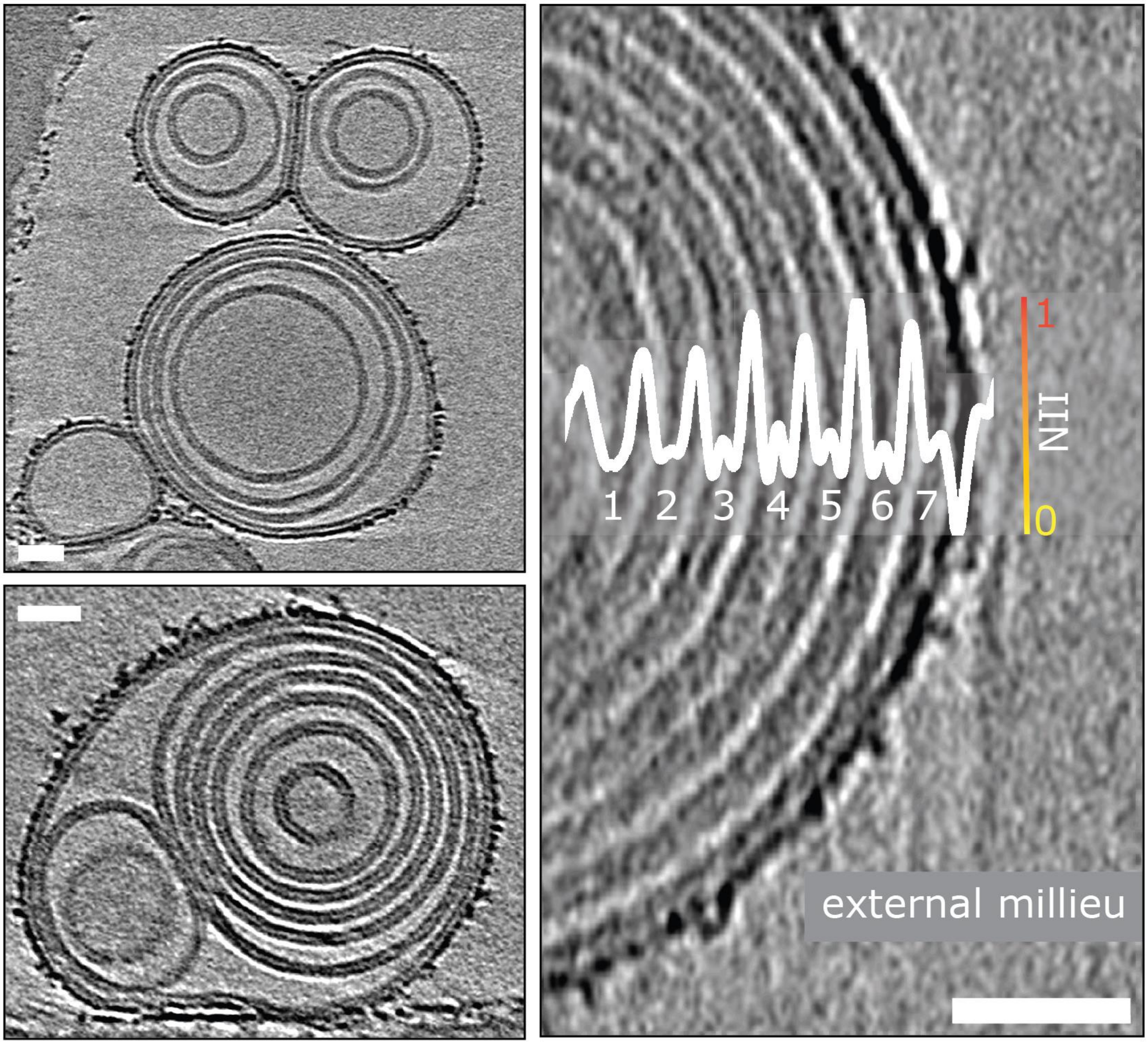


**Figure S8**: **Aβ oligomers and protofibrils do not migrate to the interior of the liposome.** Tomographic slices (7.6 nm thick) showing that only the outermost leaflet can be decorated with Aβ42 protofibrils whereas the inner layers are protected. Examples of multilamellar vesicles are shown. See Movie M5 for a tomogram of the vesicle shown in the top left panel. On the right panel seven concentric lipid bilayers (numbered) are present. The overlaid NII intensity plot (white) indicates that the outer leaflet of the outermost lipid bilayer (7) is densely packed with Aβ42 oligomers whereas bilayers (1-6) are of similar densities to the inner leaflet of bilayer (7) and devoid of Aβ42. Scale bars: 25 nm.

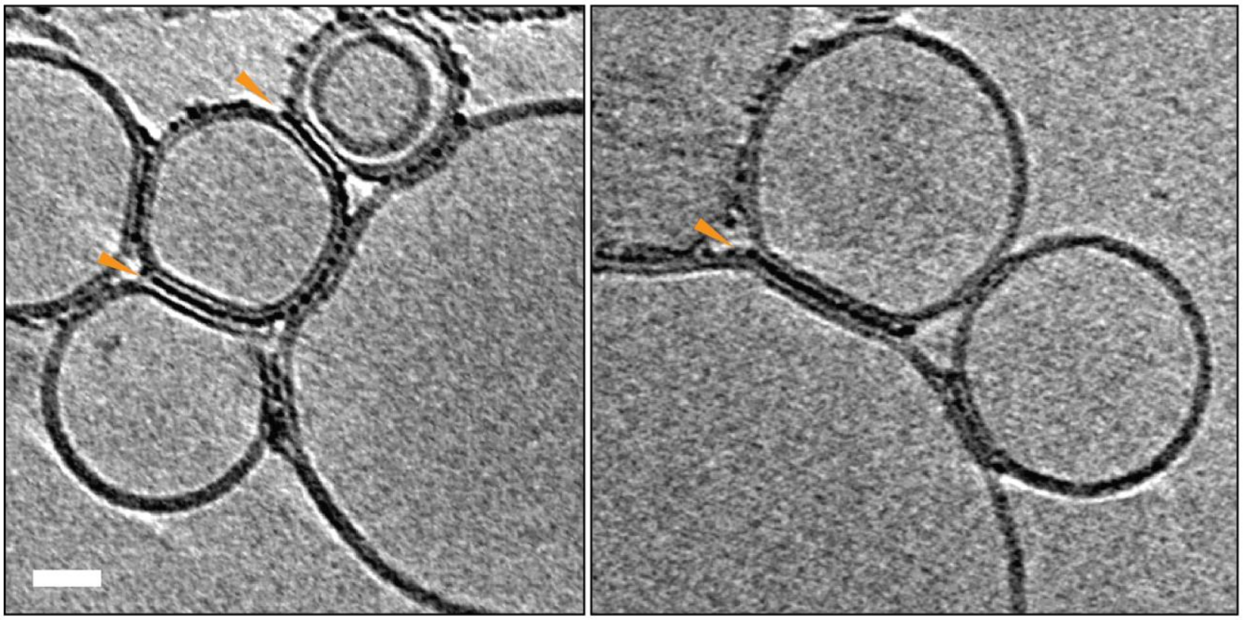

**Figure S9: Liposomes can be linked together by Aβ42 protofibrils:** For preparation with reduced levels of Aβ42 oligomer/protofibrils, Aβ42 assemblies cluster at the intervesicular space and connect the membranes of neighboring vesicles (orange arrowheads). Aβ42 is not observed elsewhere on the vesicles. The tomographic slices are 7.6 nm thick, with scale bar: 25 nm.

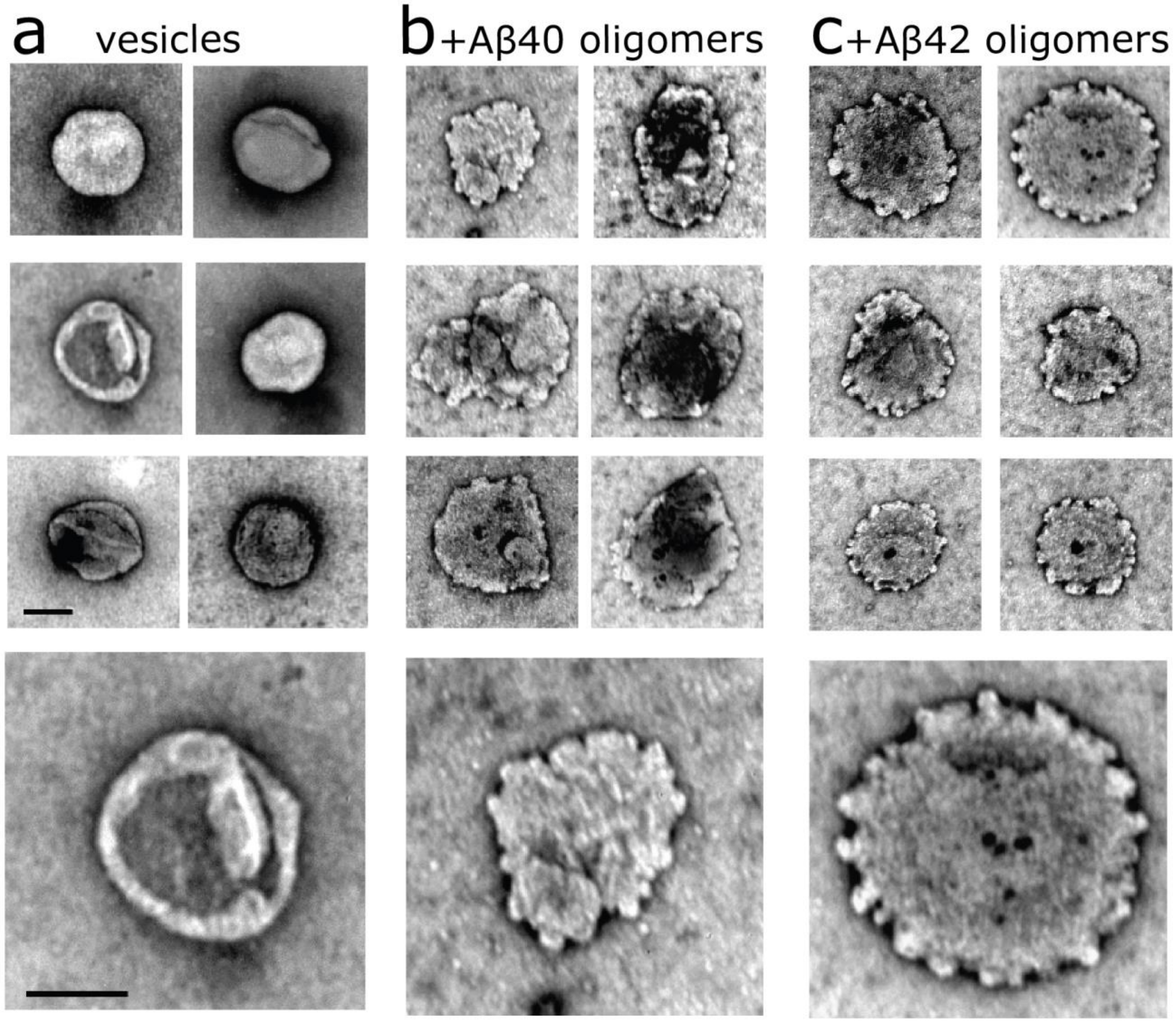


**Figure S10: Negative-stain TEM images of vesicles with Aβ oligomers**. Aβ40 and Aβ42 oligomers **(b** and **c)** (10 μM) disrupt lipid vesicles. In the presence of uranyl-acetate, negative stain, oligomer cause curvature and budding-off of membrane, while vesicles in the absence of Aβ **(a)** have relatively smooth surface. Bottom row shows enlarged images from columns above. Lipid vesicles contain, PC:Cholesterol:GM1 (68:30:2, by weight) 0.05 mg $ml^{-1}$, pH 7.4. Scale bar: 100 nm.

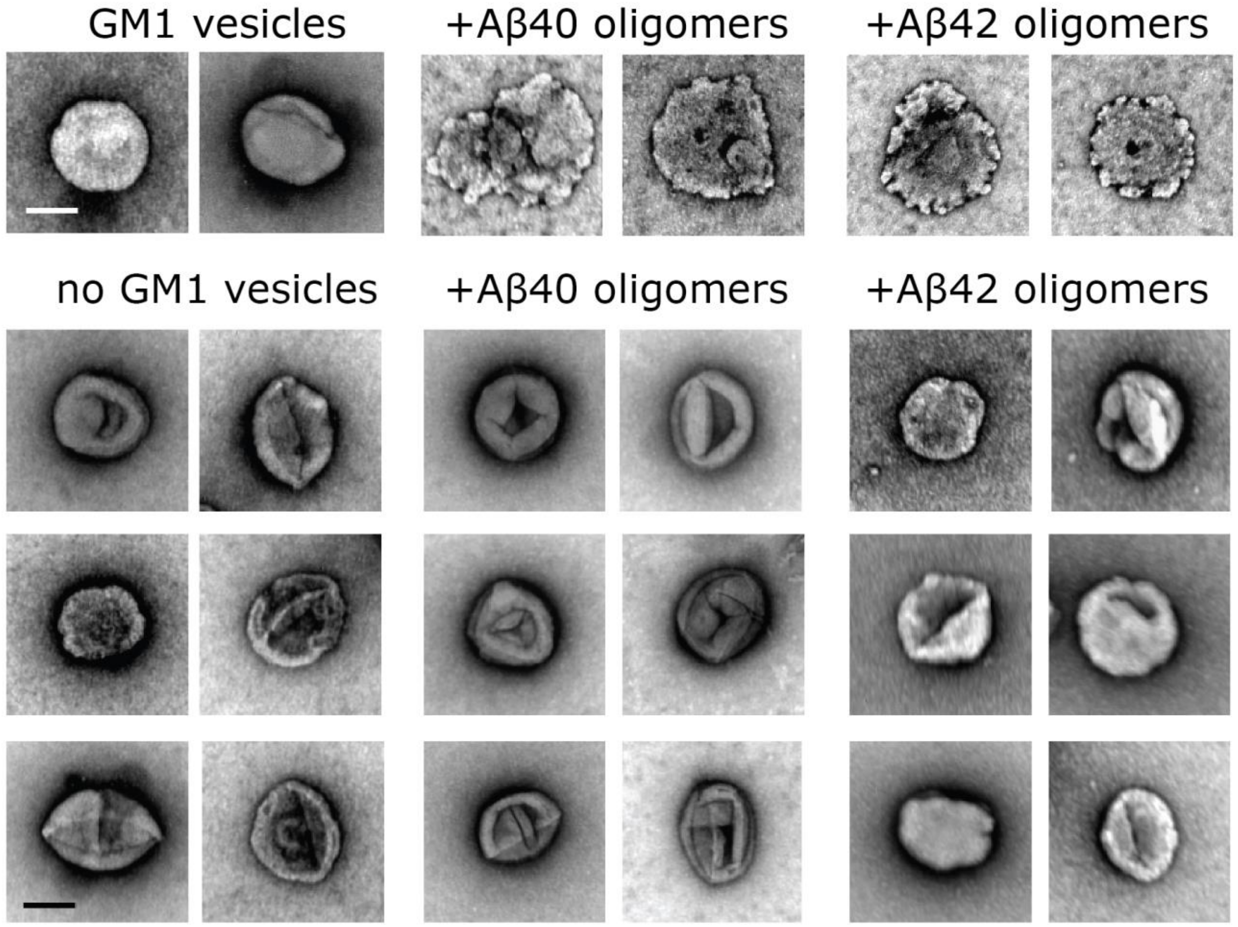


**Figure S11: Negative-stain TEM indicates GM1 is important for Aβ-lipid bilayer interactions.** Negative stain (uranyl-acetate) TEM of vesicles with Aβ oligomers. Vesicles that do not contain GM1, PC:Cholesterol only (70:30, by weight) are relatively unperturbed by Aβ40 and Aβ42 oligomers. The top row shows control the wide-spread disruption by Aβ for vesicles contain 2% by weight of GM1, as shown in supplemental Figure S10. Vesicles (0.05 mg ml$^{-1}$) were incubated with Aβ oligomer (10 μM) for 2 hrs at pH 7.4. Scale bar: 100 nm.

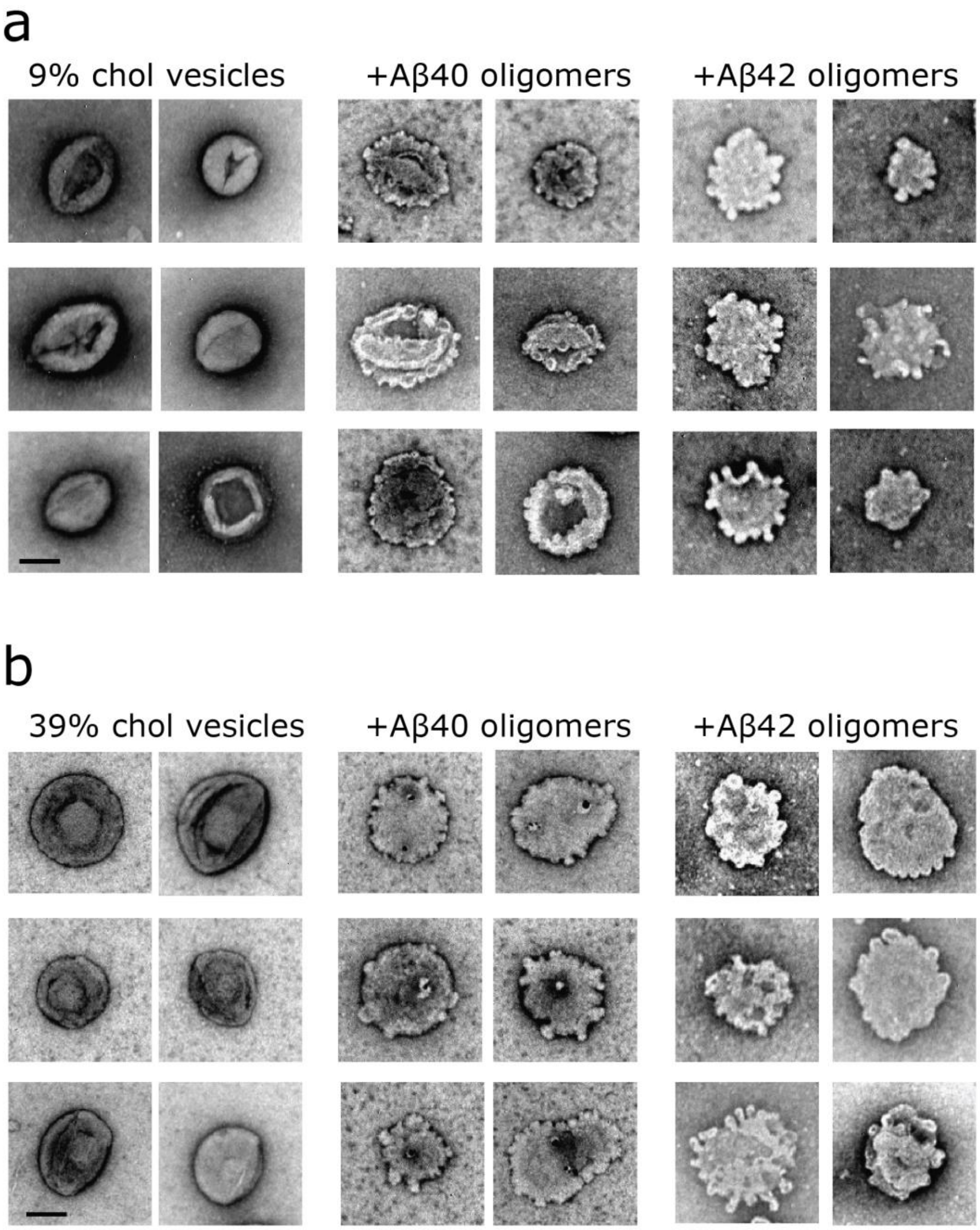


**Figure S12: Cholesterol enriched or depleted vesicles do not affect levels of Aβ oligomer induced membrane disruption.** Aβ40 and Aβ42 oligomers disrupt lipid vesicles irrespective of cholesterol levels. Negative-stain (uranyl-acetate) TEM of vesicles with and without Aβ oligomers. **a)** depleted cholesterol 9% (with 89% PC by weight). **b)** cholesterol enriched vesicles 39% (with 59% PC). All lipid vesicles contain 2% GM1. Vesicles (0.05 mg $ml^{-1}$) incubated with Aβ oligomers (10 μM) for 2 hrs at pH 7.4. Scale bar: 100 nm.

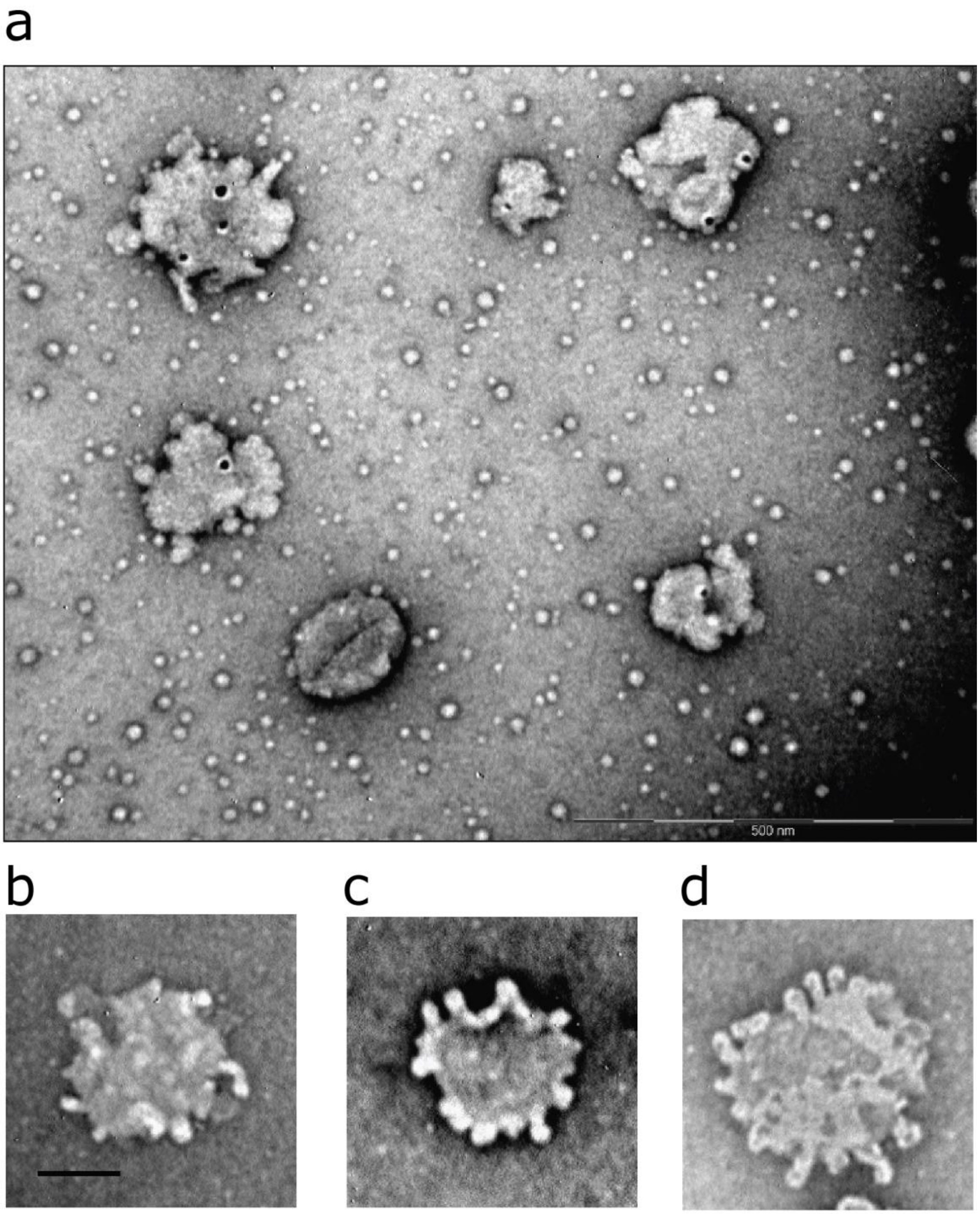


**Figure S13: Aβ oligomers induces budding-off of membrane and lipid micelle formation when in the presence of uranyl-acetate.** Negative-stain (uranyl-acetate) TEM of vesicles with Aβ oligomers. **a)** Aβ40 on 39% Cholesterol vesicles. **b)** Aβ42 on 9% Cholesterol vesicles. **c)** Aβ42 on 9% Cholesterol vesicles. **d)** Aβ42 on 39% Cholesterol vesicles (B-D are enlarged images from Supplemental Figure S12. All lipid vesicles contain 2% GM1. Vesicles (0.05 mg $ml^{-1}$) incubated with Aβ oligomers (10 μM) for 2 hrs at pH 7.4. Scale bar: 100 nm.
